# Gonadal regulation of sex-specific immunity in tuberculosis: enhanced lymphocyte function in females and dysfunctional myeloid responses in males

**DOI:** 10.64898/2026.06.11.731661

**Authors:** Manish Gupta, Nishtha Nayyar, Jessica Shen, Shichun Lun, Sabal Chaulagain, Nikita Mangla, Oscar Nino Meza, Stefanie Krug, Geetha Srikrishna, Joseph P. Hoffmann, Eileen Scully, Sabra L. Klein, William R. Bishai

**Affiliations:** Center for TB Research, Department of Medicine, Johns Hopkins School of Medicine, Baltimore, MD 21287; ICAR-NBAIR, Bengaluru, 560024, India; W. Harry Feinstone Department of Molecular Microbiology and Immunology, Johns Hopkins Bloomberg School of Public Health, Baltimore, MD 21205; Department of Medicine, Division of Infectious Diseases, Johns Hopkins School of Medicine, Baltimore, Maryland

## Abstract

Tuberculosis (TB), the world’s deadliest infection, shows higher prevalence and mortality in males than females (M/F ratio >1.7). Using Four Core Genotype (FCG) mice to decouple gonadal from chromosomal sex (XX, XY gonadal males and XX, XY gonadal females), we show that gonadal males develop accelerated disease driven by dysfunctional myeloid responses rather than impaired bacterial recognition. Both XX and XY males exhibited increased mortality, higher Mycobacterium tuberculosis (Mtb) burden, and severe lung pathology. Mechanistically, gonadal male susceptibility involved early myeloid priming, excessive neutrophil recruitment, CCR2⁺ monocyte accumulation, and hyperinflammation, with enhanced neutrophil extracellular trap (NET) formation and disorganized granulomas, implicating testes and androgens as key drivers of male susceptibility. While XY gonadal females were less susceptible, XX gonadal females showed the greatest resistance, associated with coordinated T– and B-cell responses and enhanced B-cell follicle formation. Together, these findings identify gonad-driven myeloid dysregulation as a central mechanism underlying male TB susceptibility.

## Introduction

Active tuberculosis (TB) has long been known to be more common in males than in females in high, middle, and low-income countries^1^. With the adoption of mandatory TB reporting in the early 20th century, it became clear that both the incidence and mortality of TB in males is approximately 1.7 times higher than in females, and this has been consistently observed globally by annual WHO TB reports since 1997^1^. Under-reporting of cases does not explain the difference as females have been shown to access TB care more readily and earlier than males, even in low-income settings, a factor that would lead to higher female case numbers^2^.

Beyond these epidemiologic data, natural experiments suggest that the male-female difference in TB outcome is due to biological sex differences. In the Lübeck disaster of 1929, where 251 newborns were inadvertently inoculated with virulent *Mycobacterium tuberculosis* (Mtb), mortality among boys was significantly higher (36%) than among girls (27%)^3^. Similarly, records from a Kansas institution, which used castration as a method of male behavioral control between 1871 and 1932, found that TB mortality among castrated males was significantly lower (8.1%) than that among age-matched gonadally intact men (20.6%)^4^. TB sex differences have also been observed in murine models, where female mice show better containment of Mtb proliferation than males^5, 6^.

Sex differences in infectious disease phenotypes have been shown to be modulated by both gonadal steroid hormone secretion (i.e., estrogens and progesterone in ovaries and androgens in testes) and genetic factors typically linked to X and Y chromosomes^7, 8^. Lower rates of active TB among females are first apparent following puberty, suggesting that gonadal steroids are predominant drivers^1^. Indeed, gonadal steroid concentrations vary significantly between sexes and are well known to influence the proliferation and function of virtually all immune cell types^8, 9^. In general, estrogens enhance immune function, whereas testosterone and progesterone are considered immunosuppressive^7, 10^. Female resistance to TB is sustained post-menopause, implicating additional, possible sex chromosomal factors. The X chromosome contains 800-900 genes with >60 encoding immunomodulatory proteins, including pattern recognition receptors (PRRs), transcription factors (TFs), and cytokine receptors^11^. The process of X chromosome inactivation (XCI) results in the random inactivation of genes on one of the X chromosomes in XX individuals as a mechanism of compensating for X chromosome dosage effects. Approximately 20-30% of genes on the human X chromosome and 8-15% on the mouse X chromosome escape XCI and are found at greater expression levels in cells, including immune cells, from females than males^12, 13^. The Y chromosome, in contrast, contains only 45-60 protein-coding genes, most of which are associated with testis development and spermatogenesis, and it is not believed to play a prominent role in immunity. Functional polymorphisms in several X-linked genes affect human TB susceptibility, further suggesting that sex chromosome complement may be a driver of sex differences in TB outcomes^7^.

To investigate the biologic mechanisms for the TB sex difference and whether gonadal steroids or sex chromosome complement are the primary drivers, we used the ‘four core genotype’ (FCG) mouse model^14^. Due to a translocation of the *Sry* (sex-determining region Y) locus from ChrY to Chr3, gonadal sex is determined by an autosome and independent from ChrY. Breeding C57BL/6 XY*^−Sry^* males with XX females yields XYF and XXF mice with ovaries and XXM and XYM with testes (Fig. 1a and S1a), where mice of the same gonadal sex exhibit similar serum gonadal steroid levels^15^. These four genotypes distinguish gonadal-driven phenotypes from those governed by sex chromosomal factors. Using the FCG mouse model, we observed that superior, yet balanced, T– and B-cell immunity mediates gonadal female resistance to TB, whereas gonadal males exhibit a dysfunctional myeloid response accompanied by hyperinflammation that impairs lymphocyte function. Our findings indicate that gonadal steroids are the dominant drivers of the sex differences in TB, but we find evidence that sex chromosome complement also contributes, with XX chromosome complement providing protection against TB.

**Figure 1.**
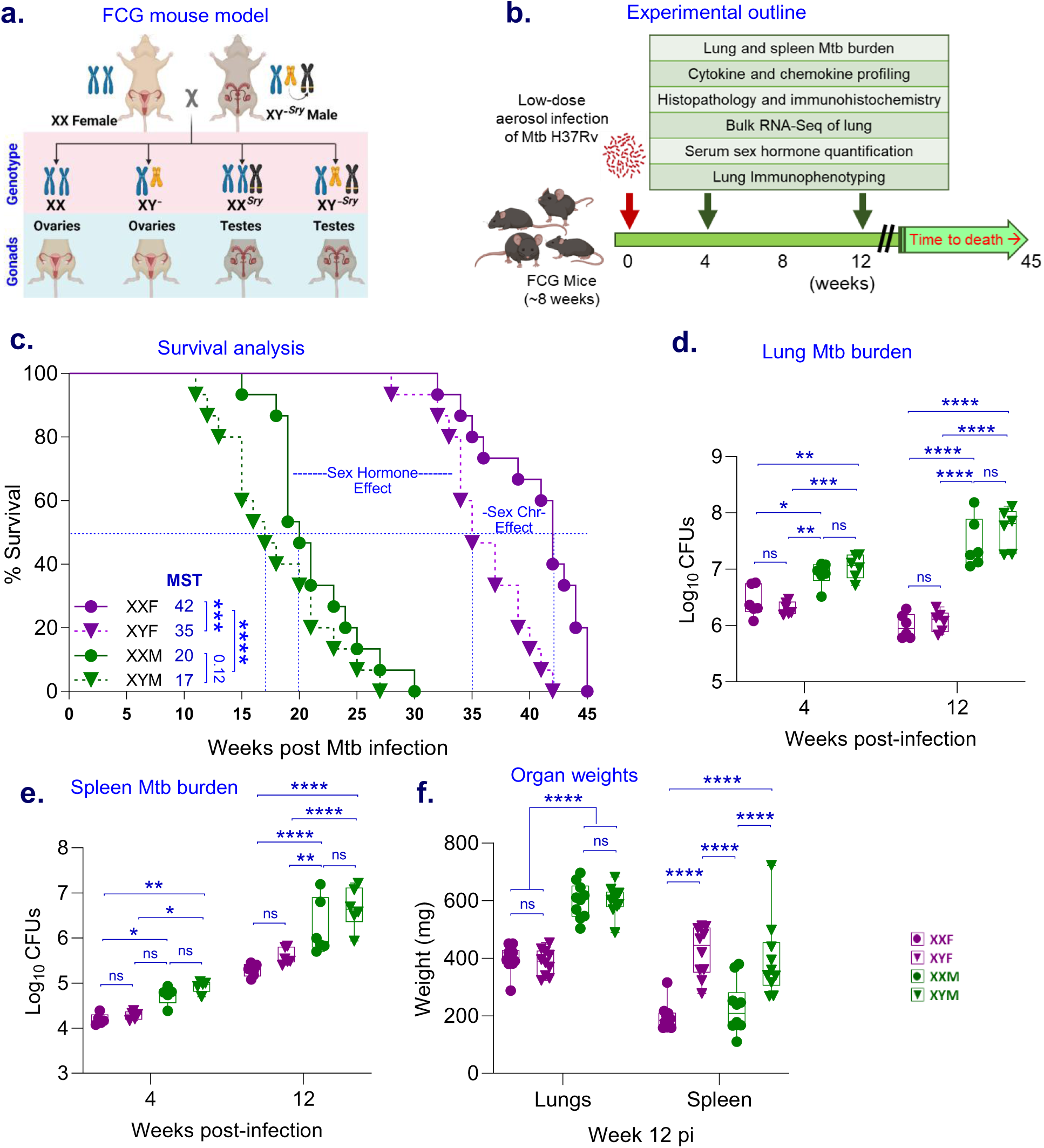
Survival kinetics and organ bacillary load in FCG mice post-Mtb infection. (**a**). The FCG mouse model. Crossing XXF with XY*^−Sry^*M produces the four core genotypes as shown– XXF and XYF with ovaries, and XXM and XYM with testes. FCG mice enable a 2 × 2 factorial comparison of sex hormones and sex chromosomes. **(b).** Schematic work plan made using BioRender. FCG mice were infected via aerosol with Mtb H37Rv (∼1.7 Log_10_ CFUs), and multiple parameters were assessed as mentioned. **(c).** Kaplan-Meier survival curve showing sex differences in TB disease progression and highlighting the influence of sex chromosomes and hormones (n = 15). Data was analyzed using the Log-rank Mantel-Cox test. Infected mice were monitored for weight loss and development of clinical symptoms (Fig. S1). At various time intervals, mice were euthanized, organs were harvested, homogenized, diluted, and plated on 7H11 selection plates. Colonies were counted after 3-4 weeks, transformed into log_10_ values, and plotted. **(d).** Lung and **(e).** spleen bacillary burden in FCG mice at 4 and 12 wpi (n = 6, except for week 4 spleen CFUs where n=5). **(f).** Gross lung and spleen weights of FCG mice (n = 10) at 12 wpi. Except for survival studies, all the experiments were performed in duplicate. Data were plotted as mean ± SEM. Statistical significance was calculated using two-way ANOVA. \**P* < 0.05, ** *P* < 0.01, *** *P* < 0.001, **** *P* < 0.0001, and ns for no significance.

## Results

### Gonadal steroids more than sex chromosome complement drives male susceptibility to TB

To evaluate TB susceptibility, FCG mice were challenged with a low-dose aerosol infection of Mtb H37Rv (∼1.7 log_10_ CFUs) and monitored for clinical indices of morbidity and mortality (Fig. 1b, S1d). Gonadal males (XXM, XYM) began losing weight at 11 weeks post-infection (wpi), with earlier symptom and higher morbidity scores, while gonadal females (XXF, XYF) continued to gain body mass through 12 wpi (Fig. S1c). Disease progression was significantly accelerated in XX and XY gonadal males compared to XX and XY gonadal females, with median times to death of 20 and 17 weeks versus 44 and 35 weeks, respectively (p < 0.0001, Fig. 1c), suggesting that testes and greater concentrations of androgens drive male hyper-susceptibility. Nevertheless, the sex chromosome complement also influenced disease progression among gonadal females only, with XYF being more susceptible to infection than XXF (p<0.001, Fig. 1c). Mtb proliferation in bone marrow-derived macrophages (BMDM) from FCG mice showed no differences suggesting that *ex vivo* macrophage function was similar across the FCG (Fig. S1g and h).

We compared Mtb burdens in the lungs and spleens of FCG mice. At 4 wpi, the lung Mtb burden was higher in gonadal males than females (p<0.05). By 12 wpi, during the chronic phase of infection, gonadal males (XXM, XYM) showed a ∼1.54 log_10_ higher lung and ∼1 log_10_ higher spleen Mtb burden than gonadal females (XXF, XYF) (p < 0.0001, Fig. 1d, S1e). Gonadal female lung CFU counts were lower at 12 wpi than at 4 wpi, indicating potent immune containment that was absent in gonadal males. Poor Mtb control in gonadal males was also apparent from the increased Mtb dissemination to the spleen in gonadal males compared to gonadal females (Fig. 1e). Greater CFU counts in gonadal males than females were accompanied by higher lung mass at 12 wpi (Fig. 1f) but not at 4 wpi (Fig. S1e). Curiously, spleen weights diverged from sex chromosome complement-driven effects, with XY showing greater splenomegaly than XX mice, independent of gonadal sex (p<0.0001, Fig. 1e). We confirmed that gonadal males of both genotypes (XXM, XYM) had equivalent testosterone levels, and gonadal females of both genotypes (XXF, XYF) had similar estradiol levels, throughout the 12 weeks of the study (Fig. S1f).

### Mtb infection causes hyperinflammation in gonadal males only

While histopathological analysis of Mtb-infected FCG mouse lungs revealed no appreciable differences at 4 wpi (Fig. S2a), striking gonadal sex-dependent differences emerged at later time points. At 12 wpi, gonadal males (XXM, XYM) exhibited extensive myeloid-dominated inflammation with poorly organized granulomas occupying >80% of lung cross-sectional area (Fig. 2a,b). In contrast, gonadal females (XXF, XYF) displayed localized, well-organized cellular granulomas covering <50% of the lung area, with prominent hematoxylin-staining lymphocytic aggregates surrounding granulomas (Fig. 2a, inset).

**Figure 2.**
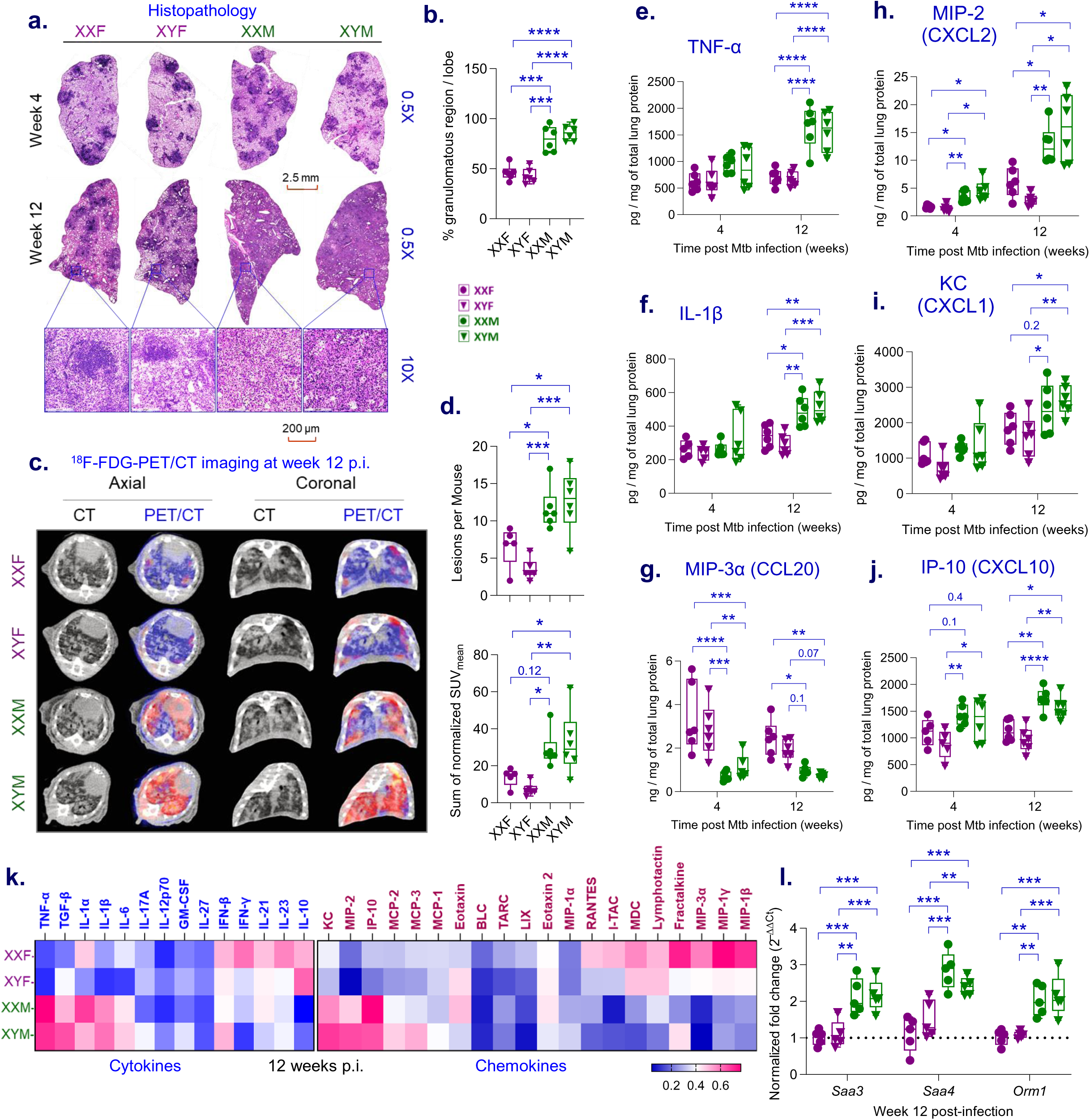
Hyperinflammation in gonadal males post-Mtb infection. Lung histopathology of Mtb-infected FCG mice was performed at 4 and 12 wpi. The lungs were formalin-fixed, sectioned, and stained with hematoxylin and eosin (H&E). **(a).** Representative H&E-stained lung sections at 0.5x magnification. Insets in week 12 sections depict lymphoid aggregates at 10x magnification. **(b).** Quantified areas of lung inflammation depicting % granulomatous region/lung section (n = 6). **(c).** ^18^F-FDG PET/CT imaging of FCG mouse lungs at 12 weeks post-infection. **(d).** Quantification includes the number of lesions per mouse and the decay– and dose-corrected standardized uptake value (SUV) of lesions, normalized to PET signals from a selected PET-blinded CT scan of non-diseased lung tissue (n = 5 for XXF and 6 for XYF, XXM, and XYM). Lung inflammatory responses in gonadal males and females were measured by quantifying cytokine and chemokine levels in lung lysates. Bar graph visualization of kinetics of pro-inflammatory cytokines– **(e).** TNF, and **(f).** IL-1β; and chemokines– **(g).** MIP-3α (CCL20), **(h).** MIP-2 (CXCL2), **(i).** KC (CXCL1), and **(j).** IP-10 (CXCL10) expression in Mtb-infected FCG lungs, at 4 and 12 wpi (n=6). Data shown as concentration normalized to total protein content. **(k).** A heat map was generated to visualize all the tested cytokine and chemokine array panels at 12 wpi by plotting C-normalized values. Min-max scaling was applied to the data for each cytokine and chemokine, adjusting the data so that the values of each variable ranged from 0 to 1.0 across all observations [C*_normalized_* =(C – C*_min_*) / (C*_max_* – C*_min_*)]. The C-normalized value for each mouse was obtained for each variable. **(l).** Normalized gene expression of various acute-phase proteins measured by qPCR at 12 wpi (n=5). Data shown as fold change relative to XXF, using the 2^−ΔΔCt^ method. Each dot represents an individual mouse. Data are represented as mean ± SEM values. Outliers are defined variable-wise by using Grubb’s test for outliers with α = 0.05. Statistical significance was calculated using two-way ANOVA. \**P* < 0.05, ** *P* < 0.01, *** *P* < 0.001 and ns for no significance.

To quantitatively assess the inflammatory burden and spatial distribution of lesions, we performed PET/CT imaging with ^18^F-FDG, which accumulates in activated neutrophils and inflammatory macrophages. At 12 wpi, gonadal males showed >2-fold increases in pulmonary lesion number and FDG standardized uptake values (SUV) relative to females (Fig. 2c–d), consistent with heightened neutrophil and inflammatory macrophage infiltration observed histologically.

Given the association between cytokine dysregulation and TB-associated lung damage, we profiled 34 cytokines and chemokines in lung homogenates by multiplex ELISA. Proinflammatory cytokines, including TNF and inflammasome-derived IL-1β were markedly elevated in gonadal males than females at 12 wpi (Fig. 2e,f,k), while IL-1α was only transiently increased in gonadal males at 4 wpi (Fig. S2b). Conversely, pulmonary concentrations of the anti-inflammatory cytokine IL-10 was significantly higher in gonadal females than males at 12 wpi (Fig. 2k, Fig. S2e), potentially contributing to their reduced inflammation. TGF-β levels were increased in gonadal males relative to XXF (Fig. 2k, Fig. S2c), consistent with fibrotic and dysregulated repair responses. IFN-γ, IL-21, and IL-23 were elevated in XXF compared to either XXM or XYM males (Fig. 2k, Fig. S2f–g), whereas GM-CSF, IFN-β, IL-6, IL-12(p70), IL-17A, and IL-27 showed no sex differences (Fig. 2k, Fig. S2h–m).

Neutrophil chemoattractants MIP-2/CXCL2 and KC/CXCL1 were significantly increased in gonadal males than females at 12 wpi (Fig. 2h,i,k), as was IP-10/CXCL10, a biomarker of TB severity (Fig. 2j,k). In contrast, MIP-3α/CCL20 was higher in females at 4 wpi (Fig. 2g,k), accompanied by elevated lymphotactin/XCL1 and MDC/CCL22 (Fig. S2o,p). Several type-1 chemokines (MIP-1α, MCP-1, RANTES at 4 wpi; MIP-1β at 12 wpi) were selectively increased in XXF versus XYF (Fig. S2r–u).

Finally, acute-phase response genes *Saa3*, *Saa4* (serum amyloid A), and *Orm1* (α1-acid glycoprotein) were upregulated >2-fold in gonadal male lungs compared with gonadal females, at 12 wpi (Fig. 2l), reflecting severe inflammation. Collectively, these data indicate that gonadal sex is the primary driver of maladaptive hyperinflammation and worsens lung pathology in both XX and XY males during chronic Mtb infection.

### Gonadal males exhibit dysfunctional myeloid response to Mtb infection

Because sex-biased gene expression is widespread across tissues, we analyzed lung transcriptomes from FCG mice at 0, 4, and 12 wpi to define the mechanisms underlying the observed hyperinflammation and poor Mtb control in gonadal males. RNA-seq confirmed consistent overexpression of Y-chromosome genes (*Eif2s3y*, *Kdm5d*, *Ddx3y*, *Uty*) in XY mice at all-time points, with *Sry* expressed exclusively in gonadal males. At both 4 and 12 wpi, genes upregulated in XXM and XYM were predominantly myeloid-derived and associated with innate immunity (Fig. 3a; Fig. S3a,b). At 4 wpi, XYM lungs showed increased expression of neutrophil chemoattractants (*Cxcl3, Cxcl7*), inflammatory mediators (*S100a8/a9*, *Il17a*), acute-phase genes (*Saa4*), and regulators of immune signaling (*Ahsg*, *Trim10*, *Fcrl5*) (Fig. 3a). By 12 wpi, XYM lungs exhibited further amplification of myeloid signatures, including neutrophil enzymes (*Mpo*, *Ngp*), PRRs (*Fpr1/2*, *Clec4e/n*), antimicrobial peptides (*Camp*), acute-phase proteins (*Saa3*, *Saa4*, *Orm1*), γδ T-cell markers (*Trdc*, *Trdv4*), and additional inflammatory genes (*Mmp12*, *Reg3g*, *Rsad2*) (Fig. 3a; Fig. S3a,b). Selected targets were validated by qPCR (Fig. S3c). Notably, calprotectin (*S100a8/a9*) and neutrophil-associated chemokines *Cxcl1*, *Cxcl2*, and *Cxcl10*—recognized biomarkers of TB severity—were markedly elevated in gonadal males, indicating early myeloid priming and recruitment preceding overt pathology.

**Figure 3.**
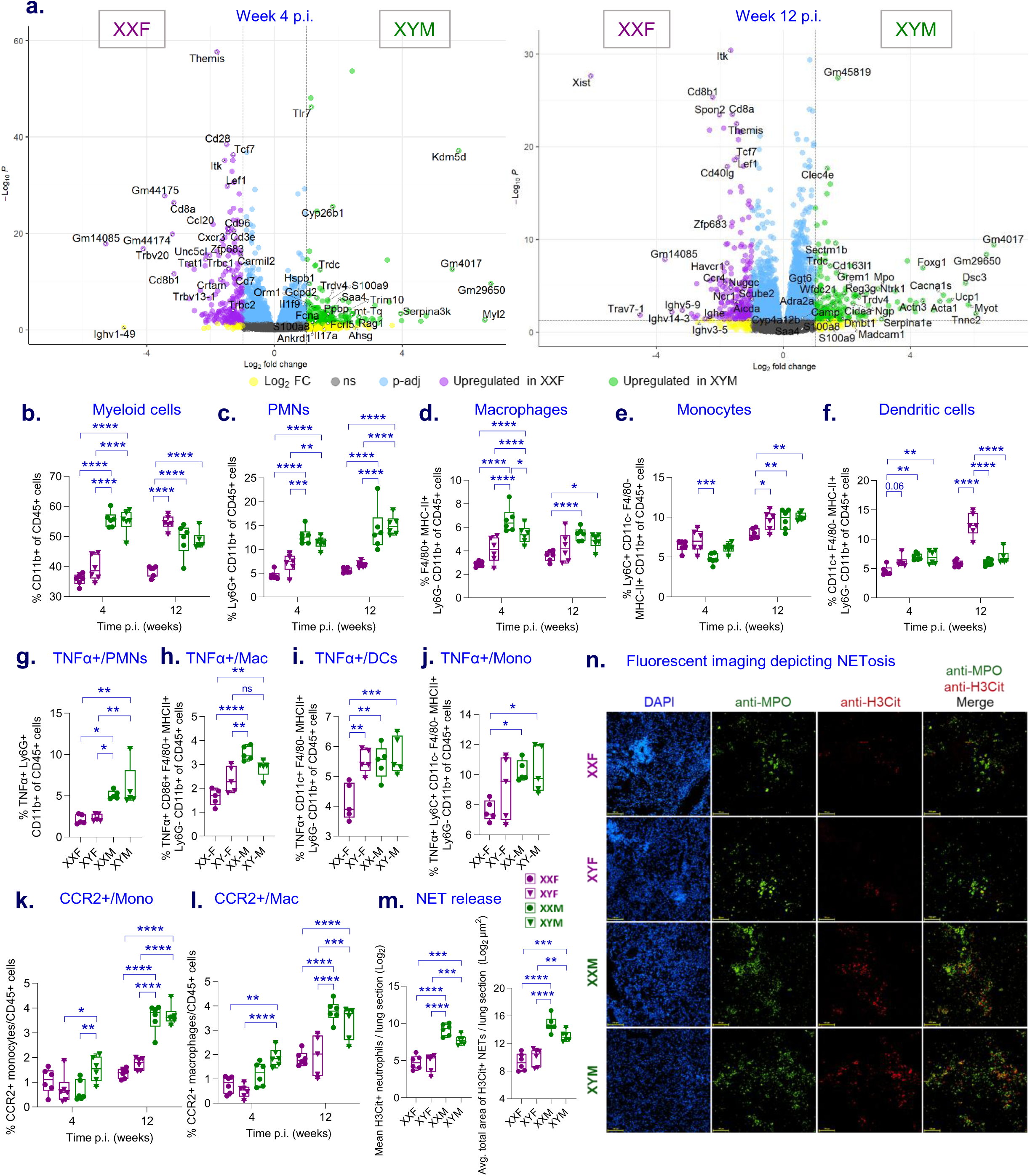
Elevated yet dysfunctional myeloid response in males after Mtb infection. (**a**). Volcano plot representing differential gene expression in XXF vs. XYM lungs after 4 and 12 weeks post-Mtb infection (n=5). The negative base-10 logarithm of *P*-adjusted values is plotted on the Y-axis, and fold change (Log_2_) is plotted on the X-axis. Purple indicates transcripts up-regulated in XXF, while green indicates transcripts up-regulated in XYM, and blue indicates no significantly differentially expressed genes (Log_2_ FC >1, *P* < 0.05). As described in the methods, FCG mice were euthanized at 4 and 12 wpi, and single-cell suspensions of their lungs were stained with appropriate antibodies and analyzed through flow cytometry. Lung immuno-phenotyping depicting the frequencies of **(b).** CD11b+ all myeloid cells, **(c).** PMNs/neutrophils (Ly6G+ CD11b+), **(d).** Macrophages (F4/80+ MHC-II+), **(e).** Monocytes (Ly6C+ F4/80– CD11c-MHC-II+) and **(f).** Dendritic cells (CD11c+ F4/80-MHC-II+) (n = 6). **(g-j).** At 4 wpi, single-cell suspensions from FCG mice lungs were stimulated for 4 h (detailed in the Methods), and TNF-α expression on different myeloid subsets was measured by multicolor flow cytometry (n=5). Figures 3b-f and 3g-j are from two different sets of experiments. Total cell frequencies of each myeloid subset related to Fig 3g-j is shown as Fig. S3e. No data for 12 wpi, due to loss of samples during processing (n = 5). **(k and l).** CCR2+ inflammatory monocytes and macrophages (n=6). Unless otherwise specified, the X-axis shows different time intervals (4 and 12 wpi), and the Y-axis displays the total frequency of the CD45+ population. Immunofluorescence staining of Mtb-infected FCG mouse lungs at 12 wpi (same sections as used in Fig. 2a), stained with fluorescently labeled antibodies to detect myeloperoxidase (MPO in green, for probing neutrophilic granules), citrullinated histone H3 (H3Cit labelling, in red), and nuclear counterstain (DAPI to stain DNA, in blue). **(m).** Bar graph displaying the mean H3Cit+ neutrophils and the average total area of H3Cit+ neutrophils per lung section, quantified using large stained areas covering the entire lung section scanned at low magnification (4x), as shown in Figure S3h, with the Fiji image analysis tool. For each mouse, 2-4 lung sections were analyzed at 4x magnification, with a total of 6 mice per FCG group. **(n).** Representative 20X scans. Each dot represents an individual mouse. Data are represented as mean ± SEM values. Outliers are defined variable-wise by using Grubb’s test for outliers with α = 0.05. Statistical significance was calculated using two-way ANOVA. \**P* < 0.05, ** *P* < 0.01, *** *P* < 0.001 and ns for no significance.

To corroborate these findings at the cellular level, lung immunophenotyping was performed. At 4 wpi, total CD11b⁺ myeloid cells were significantly increased in XXM and XYM compared to females (Fig. 3b; Fig. S3d,e). Compared with gonadal females, gonadal males exhibited ∼2-fold higher Ly6G⁺ neutrophil frequencies at both 4 and 12 wpi (Fig. 3c), alongside increased F4/80⁺ macrophages at 4 wpi (Fig. 3d). At 12 wpi, XYF mice showed elevated myeloid frequencies comparable to males but with altered composition, including increased monocytes and >2-fold expansion of pan-DCs relative to XXF (Fig. 3b,e,f).

Consistent with heightened lung TNF levels, gonadal males had significantly greater proportions of TNF^+^ neutrophils, macrophages, DCs, and monocytes (Fig. 3g–j). CCR2⁺ inflammatory monocytes and macrophages—key mediators of lung recruitment and pathology—were also enriched in XXM and XYM at 12 wpi (Fig. 3k,l).

Given the association between neutrophil activation and NETosis in TB, lung sections were analyzed by IHC for citrullinated histone H3, a key biomarker of NETs. Gonadal males exhibited increased neutrophil accumulation and aggregation (Fig. S3g,h,i) and significantly higher H3Cit⁺ NET burden and area compared to gonadal females (Fig. 3m,n). Together, these data suggest that male-biased myeloid priming, neutrophilia, CCR2-driven monocyte recruitment, and excessive NETosis collectively drive hyperinflammatory lung pathology following Mtb infection.

### Gonadal females mount greater adaptive immune responses to Mtb infection

To further define female resistance factors, we examined differentially expressed genes (DEGs) in gonadal female lungs following Mtb infection. At 4 wpi, 42 of 45 genes upregulated in XXF mice compared with XYM were associated with T cell–mediated adaptive immunity. These genes clustered into five functional groups: (i) T cell receptors and ligands (*Cd8a, Cd8b, Cxcr3, Trbc1/2, Icos, Cd40lg*, *Il27ra*), (ii) T cell development, activation, and migration (*Themis, Cd28, Cd27, Carmil2*), (iii) TCR signaling (*Trat1, Itk, Ubash3a, Prkcq*), (iv) transcription factors (*Zfp683, Lef1, Tcf7, Satb1*), and (v) cytokines and chemokines (*Ccl20, Xcl1*) (Fig. 4a). Selected targets were validated by qPCR (Fig. S4a). Comparable gonadal female-biased enrichment of T cell–associated genes was observed in XYF vs. XYM and XXF vs. XXM comparisons, contrasting sharply with the innate myeloid priming seen in males.

**Figure 4.**
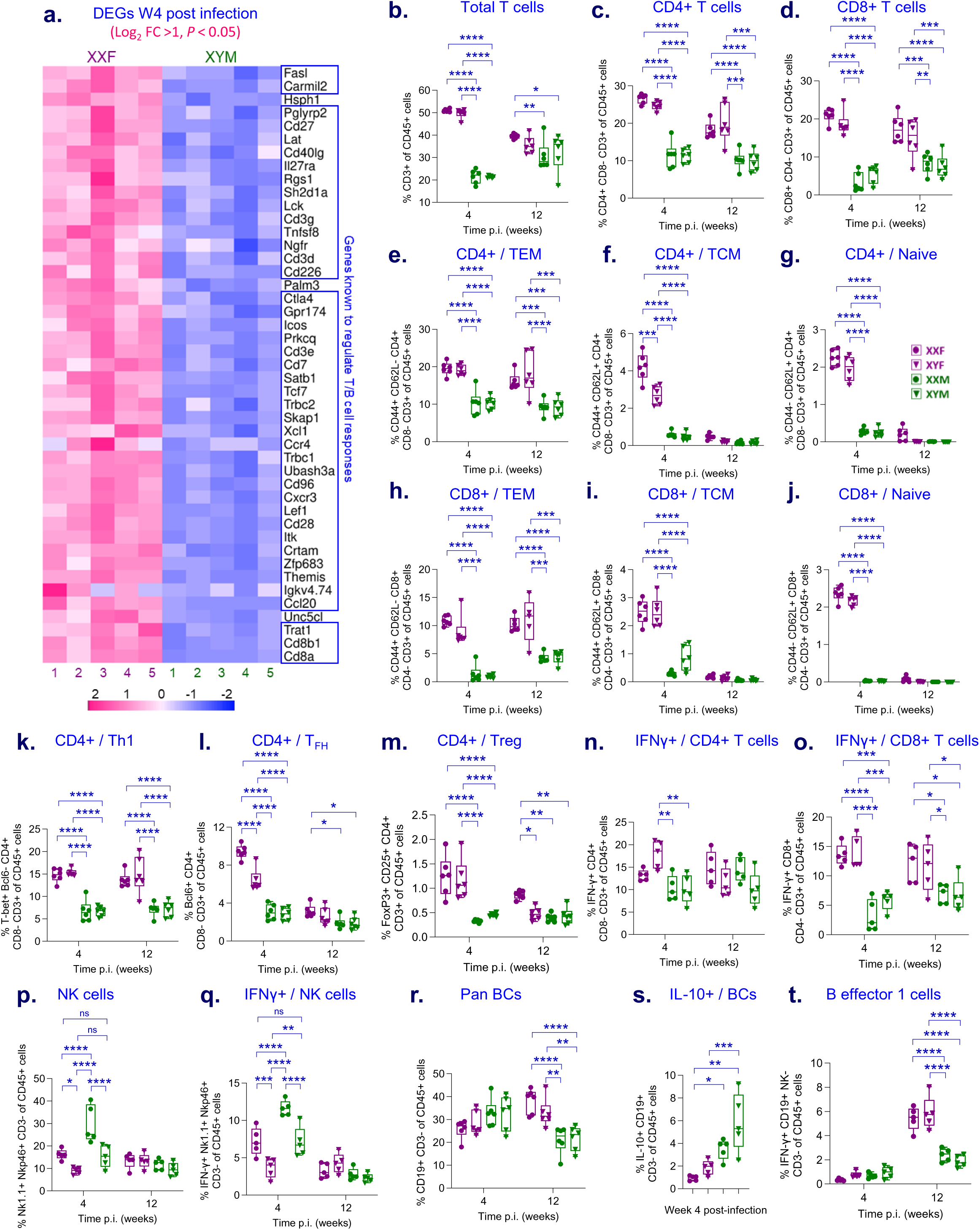
Gonadal females mount a superior T cell response to Mtb infection. (**a**). Heat map depicting transcripts of immunomodulatory genes upregulated in XXF (pink) vs. XYM (blue) at 4 wpi (n=5; Log_2_ FC >1, *P* < 0.05). Genes known to regulate T cell responses are highlighted in navy blue. As described in the methods, FCG mice were euthanized at 4 and 12 wpi, and single-cell suspensions of their lungs were stained with appropriate antibodies and analyzed through flow cytometry. Lung immunophenotyping showing– **(b).** Pan T cell frequencies, **(c).** Total CD4+ and **(d).** CD8+ T cells. **(e-j).** Frequencies of T effector memory (TEM, CD44+ CD62L-), T central memory (TCM, CD44+ CD62L+), and naïve (CD44– CD62L+) subsets based on expression of CD44 and CD62L activation markers, in CD4+ **(e-g)**, and **(h-j).** CD8+ T cells (n = 6). **(k).** Frequencies of T-bet+ CD4+ Th1 cells, **(l).** Frequencies of Bcl6+ CD4+ Tfh cells, and **(m).** Frequencies of FoxP3+ CD25+ CD4+ regulatory T cells (Tregs) (n =6). At 4 and 12 wpi, single-cell suspensions from FCG mice lungs were stimulated for 4 h (detailed in the Methods), and IFN-γ expression on different lymphoid subsets was measured by multicolor flow cytometry. **(n, o).** Frequencies of IFN-γ secreting CD4 and CD8+ T cells (n = 5). **(p).** Percent pan-natural killer cells (NK cells, Nk1.1+ Nkp46+ CD3-) and, **(q).**IFN-γ+ NK cells of all immune cells (n = 5). **(r).** Frequencies of pan-CD19+ B cells (Pan BCs, n=6), **(s).** IL-10 secreting B cells (IL-10+ BCs), and **(t).** B effector 1 cells (IFN-γ+ CD19+, n = 5). No data for IL-10+ BCs frequencies at 12 wpi, due to loss of samples during processing. Unless otherwise specified, the X-axis shows different time intervals (4 and 12 wpi), and the Y-axis displays the total frequency of the CD45+ population. Each dot represents an individual mouse. Data are represented as mean ± SEM values. Statistical significance was calculated using two-way ANOVA. \**P* < 0.05, ** *P* < 0.01, *** *P* < 0.001 and ns for no significance.

Flow cytometry confirmed enhanced adaptive immunity in gonadal females. CD4⁺ and CD8⁺ T cells were significantly more abundant in the lung of gonadal females than males at both 4 and 12 wpi (Fig. 4b–d; Fig. S4b,c). At 4 wpi, females showed marked expansion of CD4⁺ and CD8⁺ effector memory (TEM) and central memory (TCM) populations (Fig. 4e,f,h,i), as well as greater frequencies of naïve T cells (Fig. 4g,j). By 12 wpi, gonadal females maintained a steady TEM-dominant profile (Fig. 4e,h). T-bet⁺ Th1 cells were significantly enriched in gonadal females than males during both acute and chronic infection (Fig. 4k). Despite this, IFN-γ production by CD4⁺ T cells following PMA/ionomycin stimulation did not differ by either gonadal or chromosomal sex (Fig. 4n), whereas IFN-γ–producing CD8⁺ T cells were significantly more frequent in gonadal females than males (Fig. 4o). T follicular helper (Tfh) cells, critical for germinal center formation and B cell follicle development, were preferentially enriched in XXF lungs at 4 wpi compared with other FCG mice (Fig. 4l). In contrast, gonadal males showed increased Th17 differentiation (Fig. S4e) and a >50% reduction in FoxP3⁺ Tregs during acute infection as compared with gonadal females (Fig. 4m).

Sex chromosome complement influenced innate lymphocytes. For example, XX mice exhibited higher frequencies of total and IFN-γ–producing NK cells regardless of gonadal sex at 4 wpi (Fig. 4p,q). While total B cell frequencies were similar early, gonadal females showed increased B cells during chronic infection (Fig. 4r). IL-10–producing B cells were elevated in gonadal males at 4 wpi (Fig. 4s). Finally, B effector 1 cells (Be1), specialized IFN-γ–producing B lymphocytes, were expanded by >2-fold in females at 12 wpi, signifying a late Th1-boosting role (Fig. 4t). Collectively, these data indicate that female resistance to TB is driven by coordinated, hormone-influenced adaptive immune responses dominated by T cells and Be1 cells.

### Female resistance to TB linked to enrichment of B cell follicles and plasmablasts cells

To further characterize the lymphocytic aggregates surrounding granulomas in FCG mice (Fig. 2a), we analyzed lungs at 12 wpi by immunohistochemistry. Hematoxylin-stained clusters were identified as CD45R/B220⁺ B cell–rich ectopic B cell follicles (BCFs), containing follicular B cells (FoBCs), Tfh cells, and follicular dendritic cells, consistent with prior reports of sex-biased TB immunity (Fig. 5a)^16, 17^. At 12 wpi, both XXF and XYF mice exhibited significantly greater numbers and area of BCFs compared to gonadal males; however, XXF mice uniquely showed enrichment of large, well-organized follicles relative to XYF, whereas XXM and XYM mice displayed few, small, or poorly organized BCFs (Fig. 5a-c). No differences in BCF formation were observed at 4 wpi (Fig. S6a), indicating that follicle development diverges later during infection. Even at 20 wpi, gonadal males failed to develop mature BCFs, indicating a gonad-driven defect rather than delayed kinetics (Fig. 5a,b). BCF abundance in females positively correlated with lung IL-21 and IL-23 levels, cytokines known to support Tfh–B cell interactions and follicle maturation (Fig. 5d,e). These findings demonstrate that ectopic BCF formation during chronic Mtb infection is regulated by both gonadal steroids and sex chromosome complement, with the XX complement conferring the strongest phenotype.

**Figure 5.**
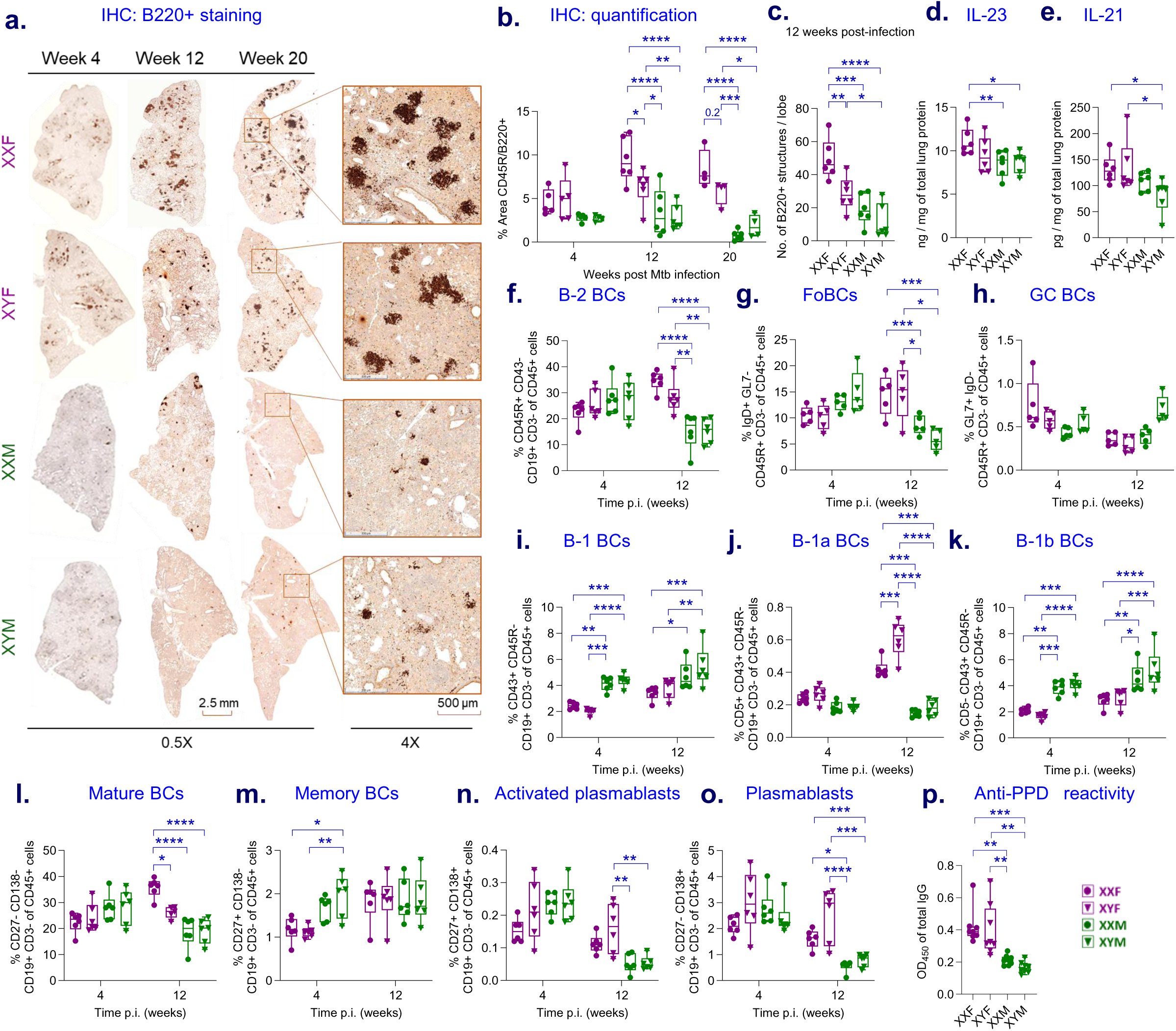
Higher BC activation and follicle formation in gonadal females. FCG mice were infected via aerosol with Mtb H37Rv as described earlier, and immunohistochemical evaluation of lung sections was performed at 4, 12, and 20 wpi. The lungs were formalin-fixed, sectioned, and stained with anti-B220/CD45R antibodies. **(a).** Representative B220-stained lung sections at 0.5x magnification. Insets in week 20 sections depict lymphoid aggregates at 4x magnification. **(b).** Quantification of total area of CD45R/B220+ structures/lung section at different time intervals and **(c).** no. of B220+ structures/lobe at 12 wpi across different FCG genotypes (n = 6). **(d).** IL-23 and **(e).** IL-21 cytokine levels in the lung lysates of Mtb-infected FCG mice at 12 wpi (n=6). Data shown as concentration normalized to total protein content. As described in the methods, FCG mice were euthanized at 4 and 12 wpi, and their lung tissues were processed into single-cell suspensions. These suspensions were stained with appropriate antibodies to identify different BC subsets, followed by flow cytometry analysis. **(f).** Frequencies of B-2 BCs (CD45R+ CD43-CD19+ CD3-, n=6). **(g).** Frequencies of follicular BCs (FoBCs, IgD+ GL7– B-2 BCs) and **(h).** Frequencies of germinal center BCs (GC BCs, GL7+ IgD-B-2 BCs) (n=5). **(i).** Expansion of innate-type B-1 BCs (CD43+ CD45R-CD19+) and frequencies of their subsets, viz., **(j)** CD5+ B-1a and **(k).** CD5-B-1b BCs (n=6). **(l-o).** Frequencies of various memory and plasma BC subsets based on expression of CD27 and CD138 (n=6). **(p).** Anti-PPD reactivity of FCG mouse sera at a 1:100 dilution, 12 weeks post-infection (n=8). Unless otherwise specified, the X-axis shows different time intervals (4 and 12 wpi), and the Y-axis displays the total frequency of the CD45+ population. Each dot represents an individual mouse. Data are represented as mean ± SEM values. Outliers are defined variable-wise by using Grubb’s test for outliers with α = 0.05. Statistical significance was calculated using two-way ANOVA. \**P* < 0.05, ** *P* < 0.01, *** *P* < 0.001 and ns for no significance.

To determine whether altered B cell differentiation caused impaired BCF formation in gonadal males, we examined lung B cell subsets by flow cytometry. Conventional B-2 B cells (CD45R⁺CD43⁻CD19⁺), which support adaptive immune memory, were significantly more abundant in XXF and XYF mice than in gonadal males at 12 wpi when they comprised >85% of lung B cells in gonadal females versus ∼70% in gonadal males (Fig. 5f). Correspondingly, FoBCs (IgD⁺GL7⁻ B-2 cells), which form the structural basis of BCFs and serve as precursors for germinal center (GC) responses, were >1.5-fold higher in XX and XY females at 12 wpi (Fig. 5g). Despite these differences, GC B cell frequencies (GL7⁺IgD⁻ B-2 cells) were comparable across sexes (Fig. 5h).

In contrast, gonadal males exhibited greater frequencies of innate-like B-1 B cells (CD43⁺CD45R⁻CD19⁺)(Fig. 5i), driven primarily by CD5⁻ B-1b subsets at both 4 and 12 wpi (Fig. 5k). Conversely, the CD5⁺ B-1a subset—associated with natural antibody production—was significantly expanded in XX and XY females at 12 wpi (Fig. 5j). Mature CD27⁻CD138⁻ B cells were also enriched in female lungs at 12 wpi (Fig. 5l). While memory B cell and activated plasmablasts or intermediate plasma cells frequencies were variable (Fig. 5m,n), short-lived antibody-secreting plasmablast cells (CD27⁻CD138⁺) were significantly increased in XXF and XYF mice at 12 wpi (Fig. 5o), accompanied by higher serum anti-PPD IgG levels (Fig. 5p). Collectively, gonadal female resistance to TB correlates with robust BCF formation, adaptive B cell differentiation, and enhanced antigen-specific antibody responses.

### Sex differences in Tfh cell help for B cell activation in TB

BCF and GC formation depend on antigen-primed B cell activation by CD4⁺ Tfh cells, creating a follicular niche that supports somatic hypermutation (SHM), class-switch recombination (CSR), and affinity maturation prior to antibody production. Whole-lung RNA-seq revealed that, compared with gonadal males, gonadal females exhibited significant upregulation of Tfh–B cell interaction signatures and BCF-associated genes at both 4 and 12 wpi (Fig. 6a). Notably, *Cd40lg* (CD40L), a key Tfh-derived signal required for B cell activation and follicle maintenance, was elevated in XXF and XYF lungs at both time points (Fig. 6a,b). Additional genes encoding Tfh costimulatory molecules (*Cd28, Icos, Fasl, Ctla4, Tnfsf8*; Fig. 6a) and B cell GC/BCF markers, including *Aicda*, *Nuggc*, and *Havcr1* (TIM-1), were similarly enriched in gonadal females compared with gonadal males (Fig. 6c-e).

**Figure 6.**
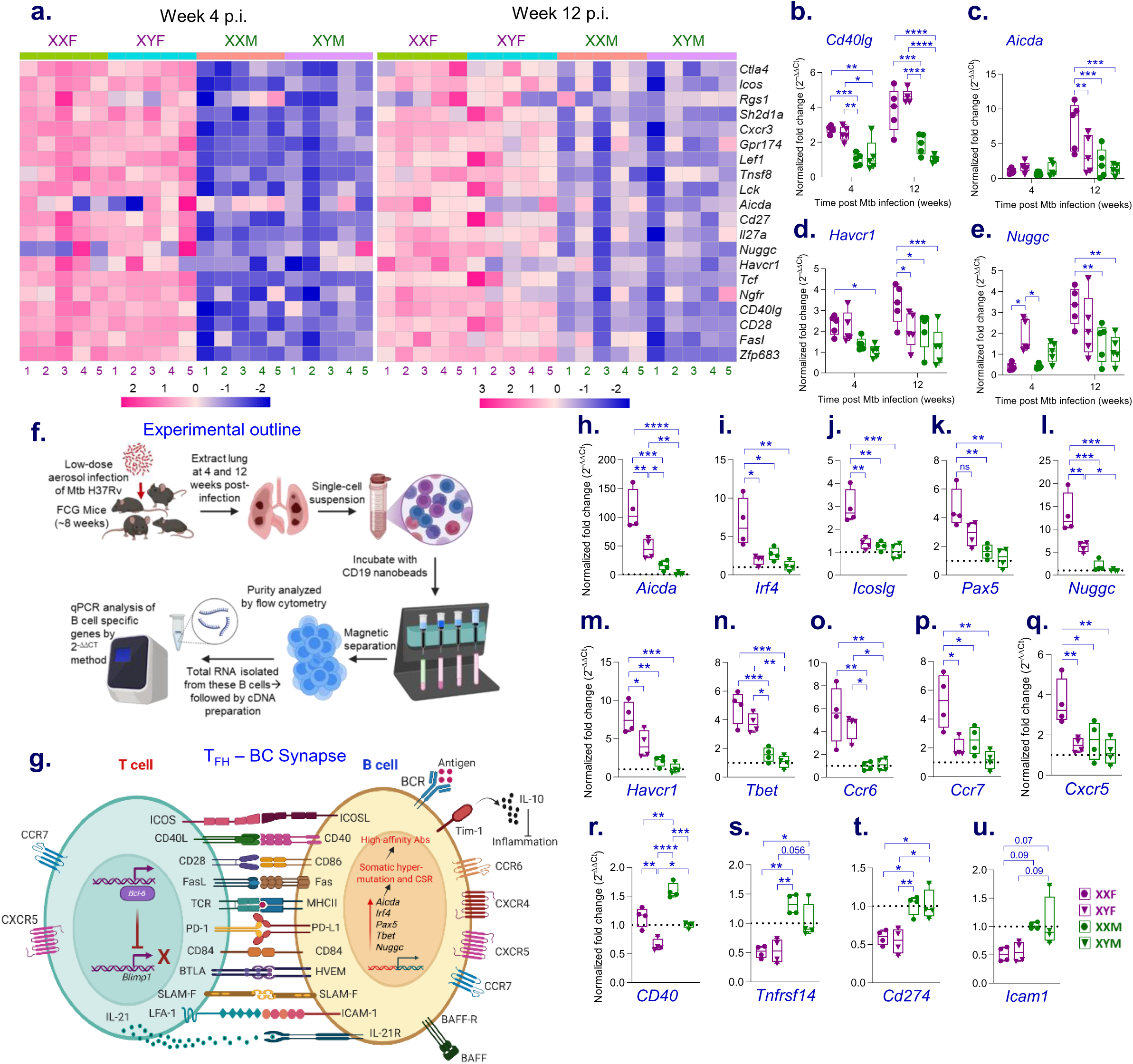
Sex differences in Tfh cell activation to BC. (**a**). A combined heat map summarizing transcripts of immunomodulatory genes known to be involved in or regulating the T-B cell synapse, across all four FCG genotypes. Upregulated transcripts are shown in pink, while downregulated ones are in blue, at 4 and 12 wpi. (n=5; Log_2_ FC >1, *P* < 0.05). **(b-e).** Validation of RNASeq data. Normalized gene expression of key genes measured by qPCR at 12 wpi (n=5). Data shown as fold change relative to XYM, using 2^−ΔΔCt^ method, for **(b)**. *Cd40lg*, **(c)**. *Aicda*, **(d)**. *Havcr1/TIM-1*, and **(e)**. *Nuggc*. **(f).** Experimental design of Mtb H37Rv aerosol infection in FCG mice, followed by B cell enrichment at various time intervals post-infection from lungs (n=4), created using BioRender.com. **(g).** Schematic diagram illustrating the T_FH_-BC synapse. **(h-u).** qPCR analysis of various genes involved in BCF formation, regulation, and maintenance. 18S rRNA normalized gene expression of various genes measured by qPCR at 12 wpi (n=4). Data shown as fold change relative to XYM (dotted line). Fold change calculated by 2^−ΔΔCt^ method. Each dot represents an individual mouse and is the average of duplicates. Data are presented as mean ± SEM values. Statistical significance was calculated using two-way ANOVA. \**P* < 0.05, ** *P* < 0.01, *** *P* < 0.001 and ns for no significance.

To directly assess differences in B cell activation, pan-CD19⁺ lung B cells were isolated (Fig. 6f), and the expression of genes involved in the Tfh-B cell synapse (Fig. 6g) was quantified by qPCR (Fig. 6h). Consistent with enhanced follicular organization, *Aicda* and *Nuggc*—both selectively expressed in GC B cells and essential for SHM and CSR—were significantly higher in XXF than XYF and gonadal males (Fig. 6h,l). IRF4, which coordinates CSR at low levels and plasmablast differentiation at higher levels, was also preferentially expressed in XXF B cells (Fig. 6j). In contrast, *Pax5* and *Tbx21* (T-bet), transcription factors involved in B cell lineage commitment and GC polarization, were expressed equivalently in XXF and XYF but at higher levels than in gonadal males (Fig. 6k,n).

Chemokine receptors that regulate B cell positioning within follicles also showed sex bias. *Ccr7* and *Cxcr5* transcripts were enriched specifically in XXF B cells, while *Ccr6* was elevated in both XXF and XYF relative to gonadal males (Fig. 6o-q). No sex differences were observed for *Ccr3* or *Cxcr4* (Fig. S7). Expression of *Icoslg* (encoding ICOSL) and *Havcr1* was significantly higher in XXF B cells than XYF and gonadal males, supporting stronger Tfh–B cell synaptic interactions (Fig. 6j,m). Unexpectedly, *Cd40* expression was highest in the XX background regardless of gonadal sex (Fig. 6r). By contrast, genes encoding *Blimp1*, *Fas*, *Cd84*, *Cd86*, and *Tnfrsf13c* (BAFF-R) were comparable across groups of FCG mice (Fig. S7).

Several negative checkpoint regulators of GC responses—including *Cd274* (PD-L1), *Tnfrsf14* (HVEM), and *Icam1*—were expressed at higher levels in gonadal male than female B cells, potentially restricting Tfh signaling and follicular maturation (Fig. 6s-u). Together, these data indicate that females, particularly XXF mice, establish a permissive Tfh–B cell activation landscape that supports functional BCF and GC development, whereas gonadal males exhibit inhibitory B cell activation signatures that limit follicular immunity during chronic Mtb infection.

## Discussion

While the drivers of the sex differences in TB morbidity and mortality are undoubtedly complex and multifactorial, our study using the FCG mouse model reveals that gonadal sex is the dominant driver of male susceptibility and female resistance. Regardless of sex chromosome complement, gonadal males (XYM and XXM) exhibited impaired Mtb containment, higher organ CFU burdens, and greater pulmonary hyperinflammation compared to XX and XY females. Sex chromosome complement exerted a secondary effect, with XXF showing greater protection against TB than XYF. Human GWAS studies support this observation as several X-linked genes are associated with sex-differential resistance to TB^7^. Sex chromosomal effects were evident only in gonadal females, likely because prolonged disease survival allowed these differences to manifest.

Sex-biased immune responses were evident during both acute and chronic TB, as reflected by differential immune cell recruitment to the lung and by cell transcriptomes. Consistent with established paradigms, the presence of testes and high concentrations of androgens promoted immunosuppression, whereas the presence of ovaries and greater concentrations of estrogens enhanced humoral immunity and optimized T–B cell balance^7, 8^. Both XXM and XYM males—with identical testosterone levels—exhibited common pathological immune features: skewing toward myelopoiesis over lymphopoiesis, sustained pulmonary hyperinflammation, excessive production of TNF, IL-1α, IL-1β, CXCL1, and CXCL2, heightened neutrophilia and NETosis, increased recruitment of inflammatory monocytes and macrophages, elevated acute-phase protein expression, and worsened lung pathology. While neutrophils contribute to early Mtb containment, excessive neutrophilia and NET formation during chronic infection promote bacterial replication, necrosis, and caseation^18, 19^. Similarly, although CCR2⁺ monocytes are required for early control, their prolonged accumulation exacerbates inflammation^20^. These maladaptive responses correlated with increased Mtb dissemination in gonadal males, whereas female lungs appeared uniquely protected from inflammation-driven tissue damage (Fig. 7).

Given the importance of cell-mediated immunity, the enrichment of CD4⁺ Th1, Tfh cells, IFN-γ⁺ CD4⁺ and CD8⁺ T cells, and immune-associated pathways in XXF and XYF likely represents a hormonally driven protective response. Sustained Th1 immunity is essential for host control of mycobacterial infection^21^. Notably, the selective expansion of IFN-γ–producing Be1 cells in gonadal females during chronic infection highlights their role as Th1 boosters^22^. Substantial evidence shows that T-bet–activated Be1 cells secrete IFN-γ in response to intracellular pathogens and promote their differentiation into antibody-secreting plasma cells^23, 24^. Reduced myeloid infiltration alone does not fully explain the moderated inflammation in female lungs. Additional protective mechanisms include elevated IL-10 levels during chronic infection^25^, increased Treg-cell expansion supporting immune homeostasis^26^, and robust BCF formation. Effective TB control depends on the coordinated transition from innate to adaptive immunity, a process shaped by complex genetic and physiological factors rather than a simple switch^8, 27^. Gonadal males—likely due to estrogen deficiency—appear unable to properly engage regulatory transcriptional programs within granulomas, leading to unchecked myelopoiesis, hyperinflammation, and impaired adaptive immunity, culminating in accelerated disease and early mortality (Fig. 7).

Our data confirm that BCF formation is a hallmark associated with female resistance to TB. Their unique presence in the lungs of gonadal females correlated with increased IL-21 expression, expansion of conventional B2 follicular B cells, elevated plasma cell frequencies, and higher anti-PPD antibody titers (Fig. 7). In contrast, gonadal males exhibited increased innate-like B1b cells, which lack immunological memory and primarily produce natural IgM^28^. The unique enrichment of IFN-γ⁺ Be1 cells in females—originating from follicular B cells via CD40–CD40L interactions^24^—supports their involvement in BCF development. Although Be1 cells have not been described in TB, they promote ectopic lymphoid follicles in *Helicobacter suis* infection^29^, supporting their potential relevance. Despite pronounced BCF differences, GC B cell frequencies were similar across sexes, possibly reflecting enhanced affinity maturation or CSR in females. While the role of BCFs in TB remains debated^30, 31^, B–cell–deficient models show minimal effects on pulmonary Mtb burden^32^ yet exhibit pronounced neutrophilia^33^. These findings suggest that ectopic BCFs may reflect an inflammation-driven lymphoid neogenesis rather than directly contributing to Mtb containment, a hypothesis that warrants further investigation.

Our findings indicate that enhanced Tfh-driven B cell activation in gonadal females, particularly XXF mice, supports robust BCF formation. In contrast, gonadal males display aberrant activation signatures coupled with elevated inhibitory checkpoint pathways like PD-L1 and HVEM, likely restricting Tfh–B cell interactions^34, 35^. This dominant inhibitory milieu provides a mechanistic explanation for defective follicular organization and diminished antibody production in males. IL-23, Tfh cells, and CD40L—critical for BCF organization^36^—were most robustly elevated in XXF mice. Importantly, the mechanistic driver of BCF formation was not strictly gonadal sex, but instead reflected coordinated regulation of genes in the B – Tfh cell molecular synapse, some of which patterned chromosome complement while others patterned gonadal sex^37, 38^. In this milieu, the expression pattern of various cell surface receptors, costimulators, TFs, and regulators of BCF/GC reactions in Mtb-infected FCGs can be clustered in three groups: a)— dependent on gonadal sex– e.g., Aicda, Tbe-t, CCR6, HVEM, PD-L1; b)— dependent on chromosomal sex– e.g., CD40; and c)— dependent on interplay between the sex steroids and chromosomes– e.g., Aicda, Ifr4, ICOSL, Pax5, Nuggc, Havcr1, CCR7, CXCR5. Study limitations include differences between murine estrous and human menstrual cycles^39, 40^, hormone dose effects^41^, and complexities of testosterone metabolism^42^. An inherent limitation of the FCG mice used here is the ∼3.2-Mb ChrX–ChrY*^Sry−^* translocation^43^, involving nine X-linked genes, including immunomodulatory loci such as *Tlr7*, *Tlr8*, and *Tmsb4x*, in an XY-biased manner, which might affect Mtb responses. However, rather than showing a survival advantage, XY-M mice were as susceptible as XX-M, and more than gonadal females, while XY-F mice remained less resistant than XX-F. Thus, translocation-driven gene overexpression is unlikely to account for the observed sex-specific differences observed in this study.

In conclusion, the FCG model has revealed several host-protective immunologic mechanisms unique to females and largely under hormonal and gonadal sex control. Future studies may enable the exploitation of these protective pathways to hone precision medicine interventions that may benefit both sexes.

## Acknowledgements

We gratefully acknowledge support from the National Institutes of Health grants R37AI167750 and P30AI16843. We thank the SKCCC OTIS Core for assistance with histology and immunohistochemistry, and the Johns Hopkins University Microbiology, Immunology, Animal Modeling & Imaging (MIAMI) Core for PET/CT acquisition and analysis. Schematic figures and the graphical abstract were created using BioRender (Academic License). Author Nishtha Nayyar was supported by funding from DST-INSPIRE Faculty Fellowship, India. We also acknowledge past and present members of the Bishai laboratory for their valuable discussions and suggestions throughout the study.

## Materials and methods

### Ethics statement

Animal studies were conducted following protocol #M022M466, approved by the Institutional Animal Care and Use Committee (IACUC) at Johns Hopkins University School of Medicine. Mice were housed in individually ventilated cages (no more than five animals per cage of the same strain) under controlled environmental conditions, including a 12-hour light/12-hour dark cycle, ambient temperatures of 20–24 °C, and relative humidity of 45–65%, with free access to food and water. At the end of the experiments, mice were humanely euthanized following IACUC guidelines to minimize pain and distress.

### Bacterial strains

The *Mycobacterium tuberculosis* H37Rv strain was sourced from the Johns Hopkins Center for Tuberculosis Research and cultured in Middlebrook 7H9 medium (Gibco) supplemented with 10% (v/v) OADC enrichment (oleic acid, albumin, dextrose, and catalase; Difco), 0.5% (v/v) glycerol, and 0.05% (v/v) Tween 80. Cultures were grown to mid-exponential phase (OD₆₀₀ ≈ 1.0), aliquoted into 1-mL volumes, and preserved at −80 °C until use.

### Animal breeding and genotyping

Prof. Arthur P. Arnold (University of California, Los Angeles) gifted breeder XY⁻ males for the FCG mouse model to Prof. Sabra L. Klein at the Johns Hopkins Bloomberg School of Public Health^14^. The FCG colonies were maintained in-house by mating XY⁻ males with wild-type C57BL/6J females (Jackson Laboratory, Strain #000664). Crosses of XY⁻(*Sry*⁺) males with XX females generate four genotypes: XX females (XXF) and XY females (XYF) with comparable adult estradiol levels, and *Sry*-containing XX (XXM) and XY (XYM) males with equivalent testosterone levels. At weaning, mice were grouped by gonadal sex (testes versus ovaries), and genotypes were confirmed by triplex polymerase chain reaction (PCR) for the presence of Sry and the Y-chromosome marker *Ssty*, as previously described^44^. Pups of the same genotype were co-housed and used for infection after 7 weeks of age.

### Bone-marrow-derived macrophage infection studies

Bone marrow–derived macrophages (BMDMs) were generated from femurs of 8-week-old FCG mice (n = 5 per genotype). Bone marrow cells were filtered through 70-µm strainers, centrifuged, and plated at 5 × 10⁶ cells/mL in BMDM medium (RPMI GlutaMAX supplemented with 10% fetal bovine serum [FBS, Gibco] and 20% L929-conditioned medium) containing 1× penicillin–streptomycin. Medium was largely refreshed every 3 days. On day 9, differentiated macrophages were harvested by gentle scraping, reseeded in 12-well plates at 2.5 × 10⁵ cells/mL in antibiotic-free BMDM medium, and stimulated overnight with recombinant murine IFN-γ (20 ng/mL) prior to infection.

BMDMs were infected with Mtb H37Rv at a multiplicity of infection (MOI) of 1:1 in BMDM medium for 4 h at 37°C and 5% CO₂. Following infection, cells were washed with pre-warmed medium to remove extracellular bacteria and treated with gentamicin (50 μg/mL) to kill residual bacilli; this was designated as time 0. Infected cells were then maintained in BMDM medium. At specified time intervals, cells were lysed with 0.1% SDS, serially diluted, and plated on Middlebrook 7H11 agar. CFUs were enumerated after 21–28 days and plotted on a log₁₀ scale. For RNA extraction, BMDMs were infected at an MOI of 1:5 for 4 h to achieve maximal infection frequency, and total RNA was extracted 6 h post-infection as described below.

### Mice aerosol infections and time to death

For all Mtb infection studies, mice were housed five per cage under biosafety level 3 (BSL-3) conditions at the Johns Hopkins School of Medicine animal facility with ad libitum access to food and water. Mice were infected at 8–10 weeks of age by aerosol exposure using a Glas-Col Inhalation Exposure System (Terre Haute, IN). Freshly thawed bacterial aliquots were diluted in sterile phosphate-buffered saline (PBS, pH 7.4) at empirically determined concentrations to achieve the target inoculum, and animals from all four genotypes were infected concurrently to minimize intergroup variability. At 24 h post-infection, 4–6 mice per run were euthanized to assess lung bacterial implantation by CFU enumeration. Animals were monitored weekly for body weight and general health. All procedures involving infected animals and infectious materials were conducted in dedicated BSL-3 facilities, and except in survival studies, moribund mice were euthanized immediately.

### Tissue collection and bacterial enumeration

Mice were euthanized at predetermined time points post-infection, and their total body weight was recorded. Terminal blood was collected from each mouse using BD Microtainer serum collection tubes, followed by the aseptic removal and weighing of lungs and spleens. The lungs were further sectioned for additional analyses: the right lobes were used for bacterial enumeration, RNA extraction, and cytokine/chemokine measurements, while the single left lobe was fixed in 10% neutral-buffered formalin for histopathology and immunohistochemistry. The weights of both the intact lung and the sampled sections were recorded to determine total bacterial burden. For CFU determination, the right lung lobes and spleens were individually homogenized in 2.5 mL of sterile PBS using CKMix50 tubes by bead-beating. Homogenates were serially diluted, and 0.5 mL aliquots were plated on Middlebrook 7H11 agar (Difco) supplemented with 10% (v/v) OADC, 0.5% (v/v) glycerol, and antibiotics (10 mg/mL cycloheximide, 50 mg/mL carbenicillin, 25 mg/mL polymyxin B, 20 mg/mL trimethoprim; Sigma-Aldrich). Plates were incubated at 37°C for 3–4 weeks before colony counting. Total CFUs per lung or spleen were calculated by adjusting for dilution, plating volume, and tissue fraction (for lungs) and expressed as log₁₀ CFUs per organ.

### Cytokine and chemokine analysis

Lung cytokine and chemokine concentrations were quantified at defined time points post-infection using pre-configured mouse LEGENDplex panels (BioLegend). Flash-frozen lung tissues were homogenized by bead beating in 2 mL Precellys CK28-R tissue homogenization tubes (Bertin Corp.) containing 1× PBS supplemented with Halt™ protease inhibitor cocktail (Thermo Fisher Scientific). Homogenates were clarified by centrifugation at 11,000 rpm for 10 min at 4 °C, and the resulting supernatants were filtered through a 0.2 μm membrane. Filtered samples were aliquoted, snap-frozen in liquid nitrogen, and stored at −80 °C until analysis. Cytokine concentrations were measured using the LEGENDplex Mouse Inflammation Panel (13-plex; Cat. #740150), while chemokines were quantified using the Mouse Proinflammatory Chemokine Panel 1 (13-plex; Cat. #741294) and Mouse Proinflammatory Chemokine Panel 2 (8-plex; Cat. #741067), in accordance with the manufacturers’ instructions. Data were acquired on a CytoFLEX flow cytometer (Beckman Coulter), and analyte concentrations were determined using the LEGENDplex™ Qognit data analysis software (BioLegend). IL-21, IL-10 and TGF-β concentrations were quantified separately using LEGEND MAX™ ELISA kits (446107, 431417, and 433007; BioLegend), following the manufacturer’s protocols. All cytokine and chemokine levels were normalized to the total protein content of the lung homogenates, as determined by a bicinchoninic acid (BCA) protein assay.

### Gonadal steroid measurement

In FCG mice, animals bearing the same type of gonad exhibit comparable circulating levels of estrogen and testosterone irrespective of sex chromosome complement (*12*). To assess the impact of Mtb infection on systemic sex steroid levels, serum concentrations were measured at 0, 4, and 12 weeks post-infection. Testosterone levels were quantified using a commercial rat/mouse Testosterone ELISA kit (IB79174; IBL America, Minneapolis, MN), and estradiol levels were measured using the EASYStep Pro Mouse Estradiol ELISA kit (RDEp-E2-Mu; Reddot Biotech Inc., Houston, TX), according to the manufacturers’ instructions.

### Lung histopathology

For histological analysis, left lung lobes were immersion-fixed in 10% neutral-buffered formalin for 72 h, paraffin-embedded, sectioned at 4 µm, and stained with hematoxylin and eosin (H&E) by the Johns Hopkins University Oncology Tissue and Imaging Service (OTIS) Core. Whole-slide images were acquired at 40× magnification using a Hamamatsu Nanozoomer S210 digital slide scanner (Hamamatsu Photonics, Shizuoka, Japan) and transferred using Concentric LS for Research (v4.4; Proscia Inc.). Regions of interest (ROIs) corresponding to inflammatory lesions were manually delineated in ImageJ (v1.54c; NIH) based on increased cellular density, disruption of normal lung architecture, and airspace obliteration. The summed ROI area for each section was normalized to the total lung area to quantify inflammatory burden. Percent lesion area was calculated as (total inflamed area × 100) / total lung area and plotted using GraphPad Prism.

### Positron emission tomography and computed tomography (PET/CT) analysis

To assess inflammatory profiles, non-invasive PET/CT imaging was employed to quantify pulmonary lesions throughout the lung and to assess their spatiotemporal distribution, which closely correlates with bacterial burden and disease severity. Briefly, FCG mice were infected with Mtb H37Rv as described above, and PET/CT imaging was performed 12 weeks post-infection using ^18^F-fluorodeoxyglucose (^18^F-FDG). Animals received a bolus intravenous injection of 7.83 ± 0.46 MBq of ^18^F-FDG (SOFIE Biosciences, Virginia, USA) via the tail vein. PET acquisition was initiated 45 min post-injection and conducted for 15 min, followed by a CT scan for anatomical reference. CT acquisition parameters were as follows: 480 projections, helical acquisition, pitch 1.0, tube voltage 50 kVp, tube current 670 µA, exposure time 300 ms, and 1:4 binning, using a nanoScan PET/CT system (Mediso, Budapest, Hungary). Co-registered PET/CT datasets were analyzed using VivoQuant 2020 (Invicro, Massachusetts, USA). Spherical volumes of interest (VOIs) were manually delineated over lesioned and non-lesioned lung regions, and radiotracer uptake was quantified. Data are presented as standardized uptake values (SUVs).

### Immunohistochemistry (IHC) analysis

Serial sections from paraffin-embedded FCG mice lung specimens were subjected to immunohistochemical staining using validated and standardized antibodies targeting B220/CD45R to identify B lymphocytes, CD11b to detect myeloid populations, and CD4 or CD8 to delineate helper and cytotoxic T lymphocytes, respectively. All staining procedures were carried out by the SKCCC OTIS at Johns Hopkins University (Baltimore, MD) on 4-μm-thick formalin-fixed, paraffin-embedded tissue sections using a Ventana Discovery Ultra autostainer (Roche Diagnostics). Sections were dewaxed and rehydrated on board, followed by heat-induced epitope retrieval using Ventana Ultra CC1 buffer (catalog #6414575001, Roche Diagnostics) at 96 °C for 64 minutes. Primary antibodies included anti-CD4 (1:1000 dilution; catalog #ab183685, Abcam), anti-CD8 (1:125 dilution; catalog #14-0195-82, Thermo Fisher Scientific), anti-CD11b (1:8000 dilution; catalog #ab133357, Abcam), and anti-CD45R/B220 (1:300 dilution; clone RA3-6B2; catalog #3088544, BD Transduction Laboratories). Primary antibodies were applied at 36 °C for 40–60 minutes, as optimized for each antibody. For rat-derived primary antibodies (CD8 and CD45R/B220), a rabbit anti-rat linker antibody (1:500 dilution; catalog #AI4001, Vector Laboratories) was applied at 36 °C for 32 minutes. Detection was performed using an anti-rabbit HQ detection system (catalog #7017936001 and #7017812001, Roche Diagnostics), followed by visualization with the ChromoMap DAB IHC Detection Kit (catalog #5266645001, Roche Diagnostics). Sections were counterstained with Mayer’s hematoxylin, dehydrated, and mounted according to standard protocols. Whole-slide scanning and imaging were conducted at the OTIS core using a Hamamatsu Nanozoomer S210 digital slide scanner (Hamamatsu Photonics, Shizuoka, Japan) at 40× resolution (0.23 microns per pixel). Image review was performed using the Concentriq pathology interface (Proscia, Philadelphia, PA) at identical settings (gamma correction 2.5, brightness 120%, and contrast 120%), and quantitative assessments were performed in ImageJ using an established analytical workflow. Tissue area measurements were generated via the color threshold function, applying uniform parameters for total pulmonary regions and DAB-reactive signal. Consistent analytical parameters were employed for all specimens, and the same threshold and detection criteria were applied uniformly to each immunohistochemical marker across slides.

### Multicolor confocal imaging

Serial 4-μm sections were prepared from paraffin-embedded lung tissues of FCG mice, mounted onto charged Superfrost Plus glass slides, and subjected to sequential deparaffinization and rehydration. Antigen unmasking was performed by heat-induced epitope retrieval using citrate buffer (Abcam) in a steamer for 30 min. Slides were rinsed in PBS, treated with TrueBlack® Lipofuscin Autofluorescence Quencher (Biotium) to reduce background signal, and blocked with 5% normal goat serum (Jackson ImmunoResearch, West Grove, PA, USA) for 1 h at room temperature. Tissue sections were incubated overnight at 4 °C with a polyclonal anti-myeloperoxidase (MPO) antibody (AF3667, R&D Systems; 1:100) and a recombinant anti-histone H3 (citrulline R17) antibody (ab219407, clone EPR20358-120, Abcam; 1:100). After PBS washes, slides were incubated for 1 h with donkey anti-goat IgG (H+L) Alexa Fluor™ Plus 488 (A32814, Invitrogen; 1:1000) and donkey anti-rabbit IgG (H+L) Alexa Fluor™ 555 (A-31572, Invitrogen; 1:1000) secondary antibodies. Nuclear counterstaining was performed with DAPI (Thermo Fisher Scientific; 1 μg/mL) for 4 min. Fluorescent images were captured using a Nikon Eclipse Ti2-E imaging system (Nikon, Minato City, Tokyo, Japan), with identical acquisition parameters applied across all experimental groups. Quantitative image analysis was performed using ImageJ/Fiji software (NIH). Uniform thresholding was applied for primary analyses, and fluorescence intensity was quantified as raw integrated density within predefined regions of interest. Lung sections from n = 6 mice per FCG group were analyzed, with at least 2 to 4 fields quantified per section, and measurements were averaged per animal before statistical analysis.

### Single cell suspension preparation

For multicolor flow cytometry, right lung lobes were excised and transferred to gentleMACS C-tubes (Miltenyi Biotec) containing the Mouse Lung Dissociation Kit enzymatic cocktail (Cat. no. 130-095-927; 1× Buffer S with Enzyme A and Enzyme D) and kept at 4 °C until processing. Lung tissue was enzymatically and mechanically dissociated into single-cell suspensions using a gentleMACS™ Dissociator with the manufacturer’s mouse lung dissociation program (37C_m_LDK_1) at 37 °C. Cell suspensions were passed through 70-µm strainers (Corning), washed with complete RPMI 1640 medium (Gibco), and centrifuged at 500 × g for 5 min at 4 °C. Red blood cells were lysed with ACK lysis buffer (Quality Biological) for 2–3 min at room temperature, after which lysis was terminated by dilution with IMDM (Gibco). Cells were pelleted at 500 × g for 5 min at 4 °C, resuspended in IMDM supplemented with 10% FBS, and counted by trypan blue exclusion using a Countess automated cell counter (Thermo Fisher Scientific).

### Multicolor flow cytometry analysis

Lung-derived single-cell suspensions were distributed into U-bottom 96-well plates (Thermo Scientific™ Nunc) at a density of 2 × 10^6^ viable cells per well and pelleted by centrifugation at 450 × g for 5 min at 4 °C. Cells were resuspended in IMDM complete medium supplemented with GolgiPlug™ (BD Biosciences; 1:1000, v/v) and Cell Stimulation Cocktail (eBioscience; 1×) and incubated for 4 h at 37 °C in a humidified atmosphere containing 5% CO₂. Following stimulation, cells were washed twice with PBS and stained with Zombie Red™ Fixable Viability Dye (BioLegend; 1:2000, v/v) for 15 min at room temperature in the dark, followed by washing. Fc receptor–mediated nonspecific binding was blocked by incubation with TruStain FcX™ PLUS (anti-mouse CD16/CD32; Cat. #156604, BioLegend) diluted in Cell Staining Buffer (CSB; 420201, BioLegend) for 20 min at room temperature. For surface marker staining, cells were incubated with antibody cocktails prepared in CSB for 30 min at room temperature in the dark. Cells were subsequently fixed using Cytofix/Cytoperm™ (Cat. #554714, BD Biosciences) for intracellular cytokine staining (IFN-γ, TNFα, and IL-10) or the FOXP3 Fix/Perm Buffer Set (Cat. #421403, BioLegend) for transcription factor analysis, including FoxP3, T-bet, RORγt, Bcl6, and GATA3. Fixation was performed for at least 1h at 4°C, followed by permeabilization using the corresponding wash buffers. Intracellular or intranuclear staining was conducted at 4 °C in the respective permeabilization buffer according to the manufacturer’s protocols. After three washes, cells were resuspended in Cyto-Last™ Buffer (422501, BioLegend) and stored at 4 °C until acquisition. Prior to analysis, samples were filtered through strainer-capped FACS tubes (352235, BD). The following anti-mouse flow cytometry antibodies were utilized, Brilliant Violet 510™ CD45 (103138, BioLegend), Pacific Blue™ CD3 (100214, BioLegend), PerCP CD4 (100431, BioLegend), Alexa Fluor^®^ 700 CD8a (100730, BioLegend), Brilliant Violet 605™ CD19 (115540, BioLegend), APC CD25 (102012, BioLegend), KIRAVIA Blue 520™ NK-1.1 (156521, BioLegend), PE/Cyanine7 CD335/NKp46 (137617, BioLegend), PE FOXP3 (320008, BioLegend), PerCP/Cy5.5 CD3 (100218, BioLegend), APC/Fire 750 CD4 (100460, BioLegend), KIRAVIA Blue 520 CD62L (104464, BioLegend), Brilliant Violet 650 CD44 (103049, BioLegend), Alexa Fluor 647 Bcl-6 (648306, BioLegend), Brilliant Violet 421 GATA3 (653814, BioLegend), PE/Cyanine7 T-bet (644824, BioLegend), PE RORγt (12-6981-80, Thermo Fisher Scientific), Alexa Fluor^®^ 700 CD11b (101222, BioLegend), APC/Fire™ 750 F4/80 (123151, BioLegend), Brilliant Violet 421 CD163 (155309, BioLegend), Brilliant Violet 650™ CD86 (105035, BioLegend), BUV563 Ly-6G (612921, BD Biosciences), Brilliant Violet 785 Ly-6C (128015, BioLegend), KIRAVIA Blue 520™ CD11c (117363, BioLegend), Spark UV387 anti-mouse I-A/I-E (107669, BioLegend), PE CD192/CCR2 (150609, BioLegend), Alexa Fluor 700 CD3 (100216, BioLegend), Brilliant Violet 421 CD45R/B220 (103240, BioLegend), APC/Fire 750 CD19 (115558, BioLegend), Alexa Fluor 488 CD5 (100612, BioLegend), APC CD43 (143208, BioLegend), Brilliant Violet 650 CD138 (142518, BioLegend), PE/Cyanine7 CD27 (124216, BioLegend), PerCP/Cyanine5.5 IgD (405709, BioLegend), and PE GL7 (144608, BioLegend). All antibodies were titrated to determine optimal staining concentrations that maximized specificity while minimizing spectral spillover. Data were acquired on BD LSR II or LSRFortessa™ flow cytometers (BD Biosciences) and analyzed using FlowJo software (v10.10; Tree Star). Compensation matrices and gating strategies were established using single-stain and fluorescence-minus-one (FMO) controls, with lung-derived single-cell suspensions used for both compensation and control samples rather than beads. Only antibodies validated for specificity by the manufacturers were employed. Gating strategies are provided in Supplementary Fig. S3f, S5a-c, and S6b.

### Antibody titers

Tuberculin purified protein derivative (PPD)-specific IgG responses were quantified by indirect enzyme-linked immunosorbent assay (ELISA). High-binding 96-well plates (Nunc MaxiSorp) were coated overnight at 4 °C with capture PPD (1 µg/well; Mantoux, TUBERSOL, 49281-752-22; Sanofi) in PBS and subsequently blocked with 2% bovine serum albumin (BSA) in PBS for 2 h at room temperature. Serial dilutions of serum samples (1:10^2^) from immunized mice were applied in triplicate and incubated for 1 h at room temperature. Plates were washed four times with PBS containing 0.05% Tween-20 (PBS-T) before incubation with horseradish peroxidase (HRP)–conjugated goat anti-mouse secondary antibodies (1:10,000; 31430, Thermo Fisher Scientific) at 37 °C for 1 h. Following additional washes, color was developed with the OptEIA TMB substrate (BD Biosciences), and the reaction was stopped with 1 N HCl. Absorbance was measured at 450 nm with wavelength correction at 570 nm where applicable, after background subtraction using uncoated wells.

### B cell enrichment

After preparing single-cell suspensions, B cells were enriched using MojoSort™ Mouse CD19 Nanobeads (BioLegend) following the manufacturer’s instructions. Briefly, cell suspensions prepared without erythrocyte lysis were filtered through 70-µm strainers and resuspended in ice-cold MojoSort™ Buffer. Cells were adjusted to 1 × 10⁸ cells/mL, incubated on ice with antibody-conjugated CD19 Nanobeads for 20-30 minutes, and then subjected to magnetic separation. The unlabeled cells were discarded, and the labeled cells were gently resuspended in MojoSort™ Buffer and underwent two additional rounds of magnetic separation to improve purity. The purified B cells were immediately used for total RNA isolation.

### RNA extraction and quantitative real-time PCR (qPCR)

Flash-frozen Mtb-infected FCG mouse lung tissues were homogenized in QIAzol, and total RNA, including small RNA species, was isolated using the miRNeasy Mini Kit (Qiagen) according to the manufacturer’s instructions. Residual genomic DNA was removed by DNase I treatment (Promega). Total cDNA was synthesized from purified RNA using the High-Capacity cDNA Reverse Transcription Kit (Thermo Fisher Scientific). Quantitative real-time PCR was performed using PowerTrack™ SYBR Green Master Mix (A46109, Thermo Fisher Scientific), either on a StepOne Plus or QuantStudio 3 Real-Time PCR System (Applied Biosystems). Primer sequences (Supplementary Table S1) were designed using the GenScript Real-Time PCR Primer Design Tool and synthesized by Integrated DNA Technologies. Data obtained from the StepOne Plus real-time PCR System were analyzed using StepOne Software v2.3 (Applied Biosystems), while those obtained from the QuantStudio 3 were analyzed using QuantStudio^TM^ Design and Analysis Software v1.5.2. Cycle threshold (CT) values were normalized to 18S rRNA to calculate ΔCT values. Based on gene up– or down-regulation patterns in gonadal males or females, ΔΔCT values were determined relative to the mean ΔCT of XYM or XXF controls, as appropriate. Relative gene expression was calculated using the 2^−ΔΔCT^ method and expressed as fold change compared with the corresponding XYM or XXF group, as indicated in the figure legends.

### Bulk RNA sequencing (RNA-seq) and analysis

Total RNA samples were shipped on dry ice to GENEWIZ (Azenta Life Sciences, South Plainfield, NJ, USA) for quality control and sequencing. RNA concentration was quantified using the Qubit RNA HS Assay Kit (Thermo Fisher Scientific, Waltham, MA), and RNA integrity was assessed with an Agilent 2100 Bioanalyzer (Agilent Technologies, Santa Clara, CA). RNA-seq libraries were prepared using a ribosomal RNA depletion strategy. Briefly, RNA was fragmented and reverse-transcribed using random primers, followed by end repair, 5′ phosphorylation, 3′ dA tailing, and adapter ligation. Libraries were amplified by PCR, and library size distribution and concentration were evaluated using the Agilent 2100 Bioanalyzer. Paired-end sequencing (2 × 150 bp) was performed on an Illumina NovaSeq platform, generating approximately 40 million read pairs per sample. Raw sequencing reads were trimmed to remove adapter sequences and low-quality bases using Trimmomatic (v0.36)^45^. High-quality reads were aligned to the Mus musculus reference genome (30-781178413_mouse_TB; Ensembl) using STAR (v2.5.2b)^46^. Gene-level read counts were generated with featureCounts from the Subread package (v1.5.2), retaining only uniquely mapped reads overlapping annotated exons. Differential gene expression analysis was conducted using DESeq2^47^. Pairwise comparisons between experimental groups were performed using the Wald test to estimate log₂ fold changes and associated p values. Genes with an adjusted p value < 0.05 and an absolute log₂ fold change > 1 were considered differentially expressed. Heatmaps were generated using the pheatmap package in R, and volcano plots were produced using the EnhancedVolcano package.

## Statistical analysis

All statistical analyses were performed using GraphPad Prism (v10; GraphPad Software, Boston, MA). Data were primarily analyzed by comparing values across time points in FCG mice using two-way analysis of variance (ANOVA) followed by Tukey’s multiple-comparisons test.

Alternative statistical approaches, when applied, are specified in the corresponding figure legends. No statistical methods were used to predetermine sample size. Experiments were randomized, and all data collected are reported in the manuscript. Colony-forming unit (CFU) data were log₁₀-transformed prior to analysis. A two-sided p-value < 0.05 was considered statistically significant. Data are presented as mean ± SEM. In animal experiments, infrequent premature mortality led to reduced sample sizes in certain groups. No data were excluded from the analyses. Following aerosol infection, animals were randomly allocated to experimental groups. Investigators were not blinded; however, they were aware of group assignments during experimental procedures and outcome assessment.

**Table S1:**
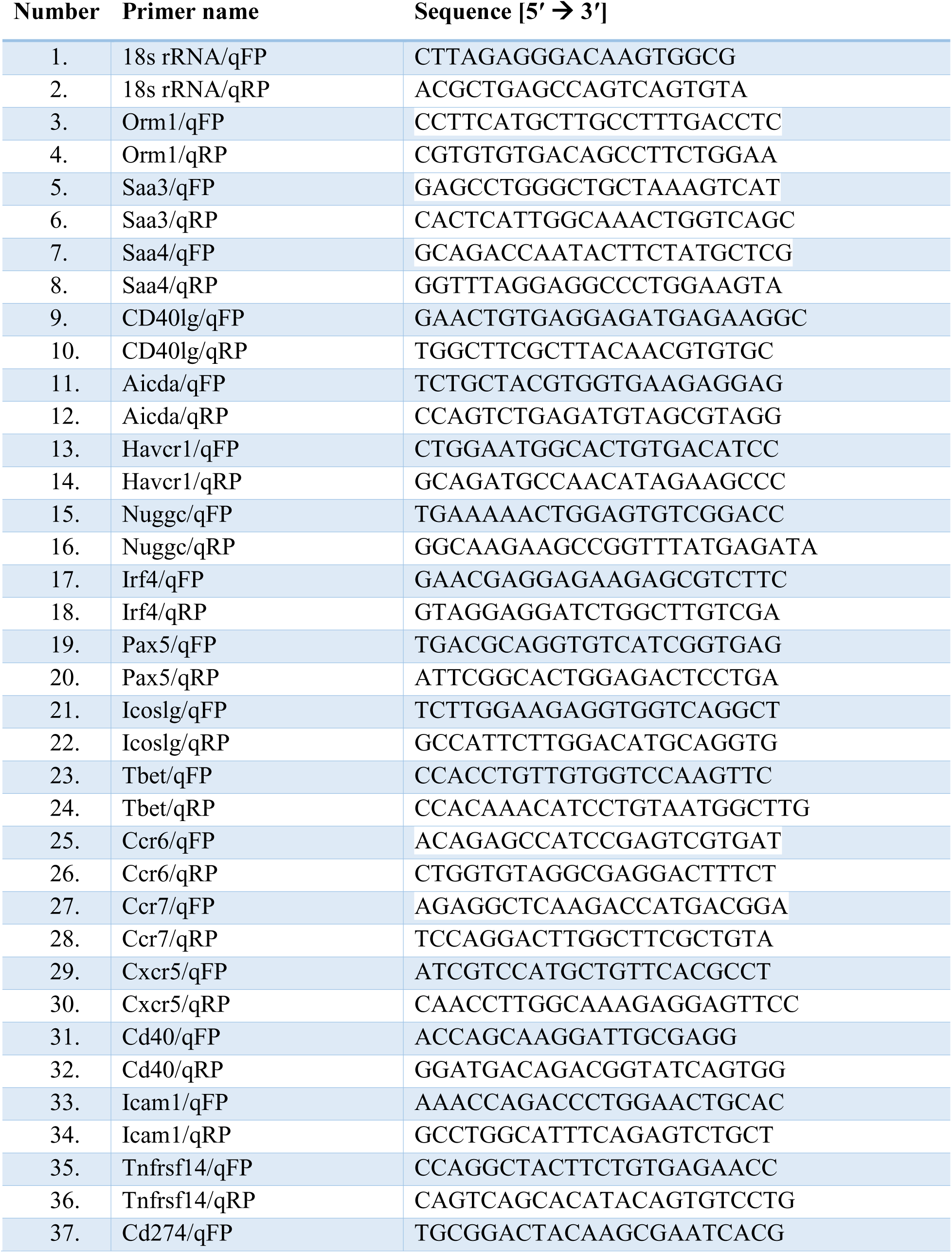
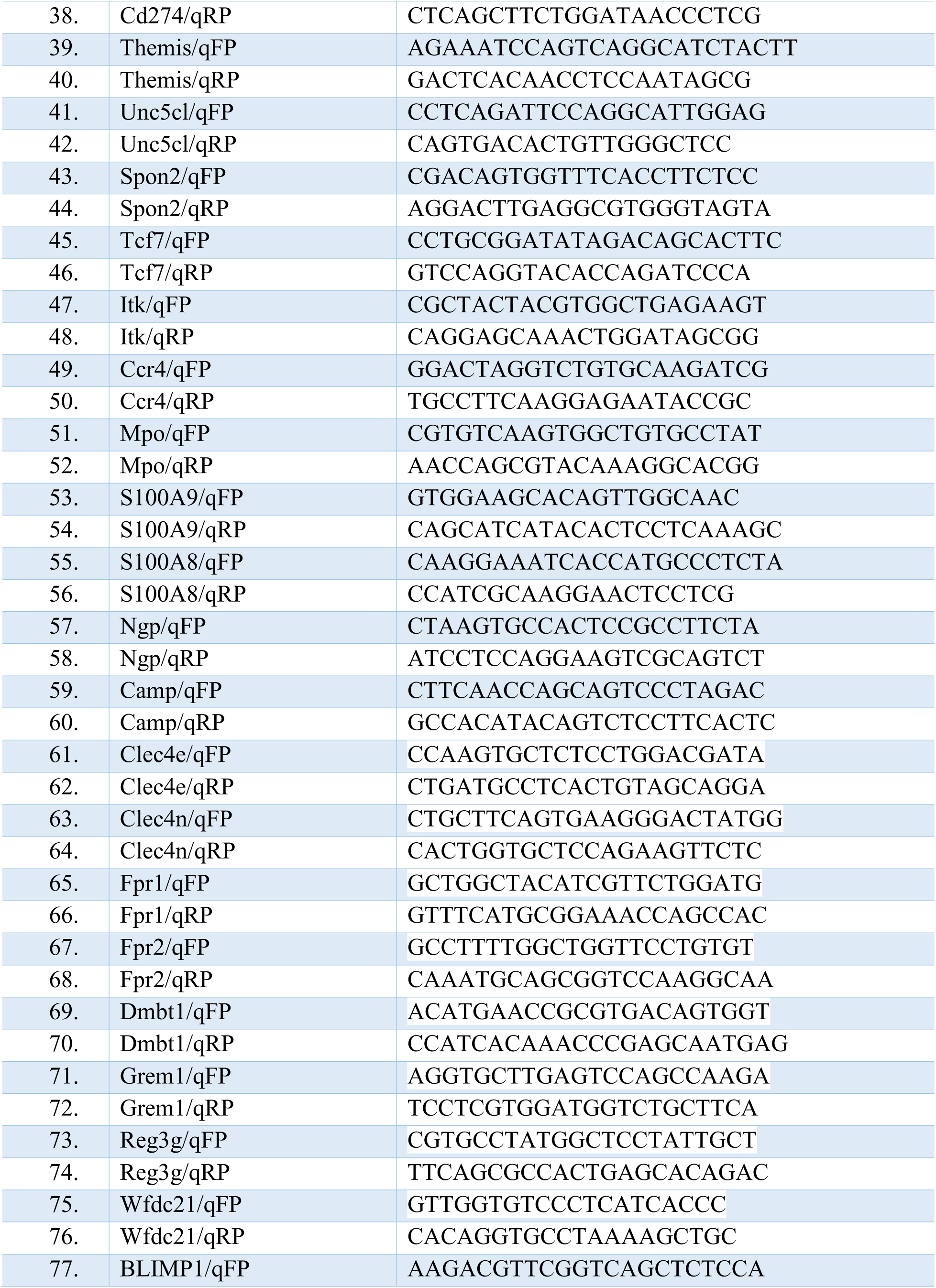
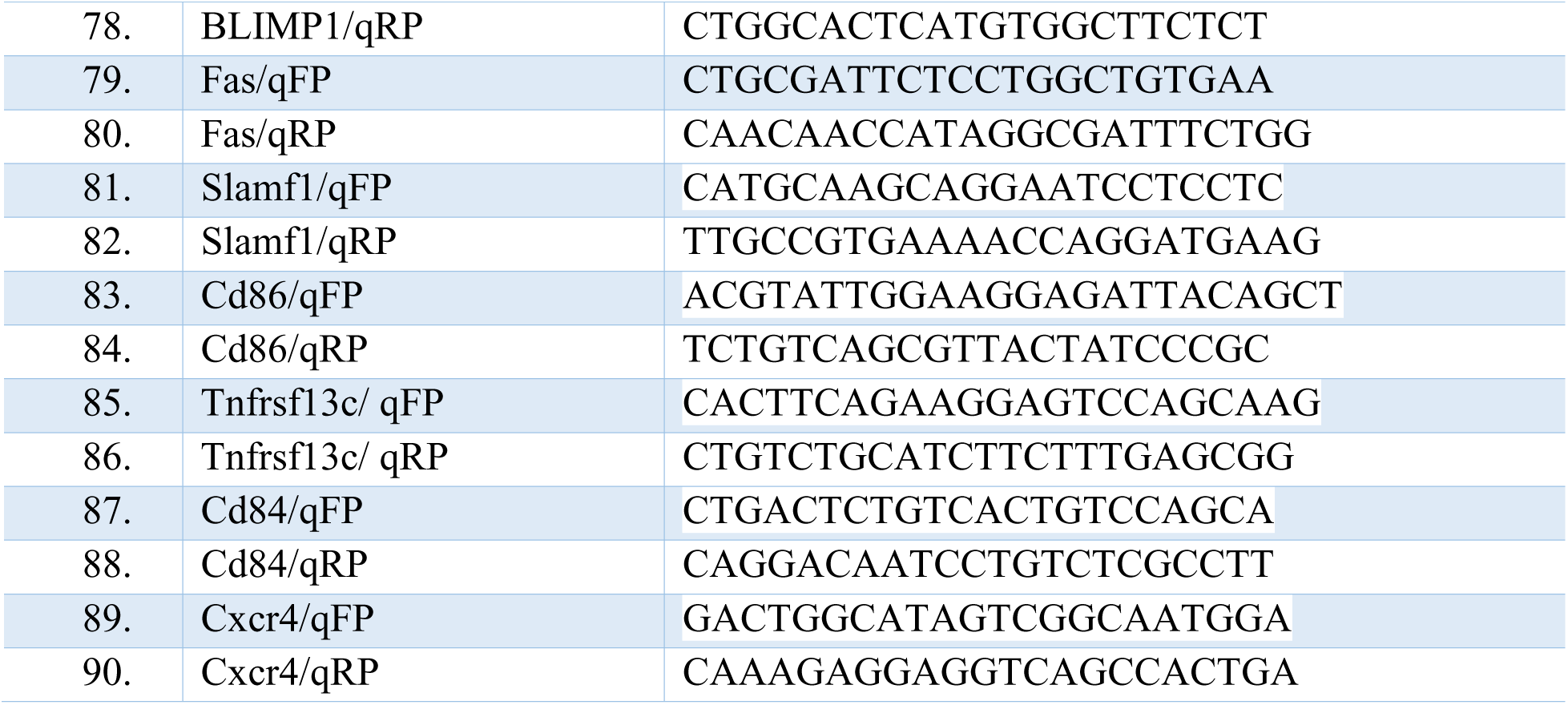
List of qPCR primers used.

---

**Figure 7.**
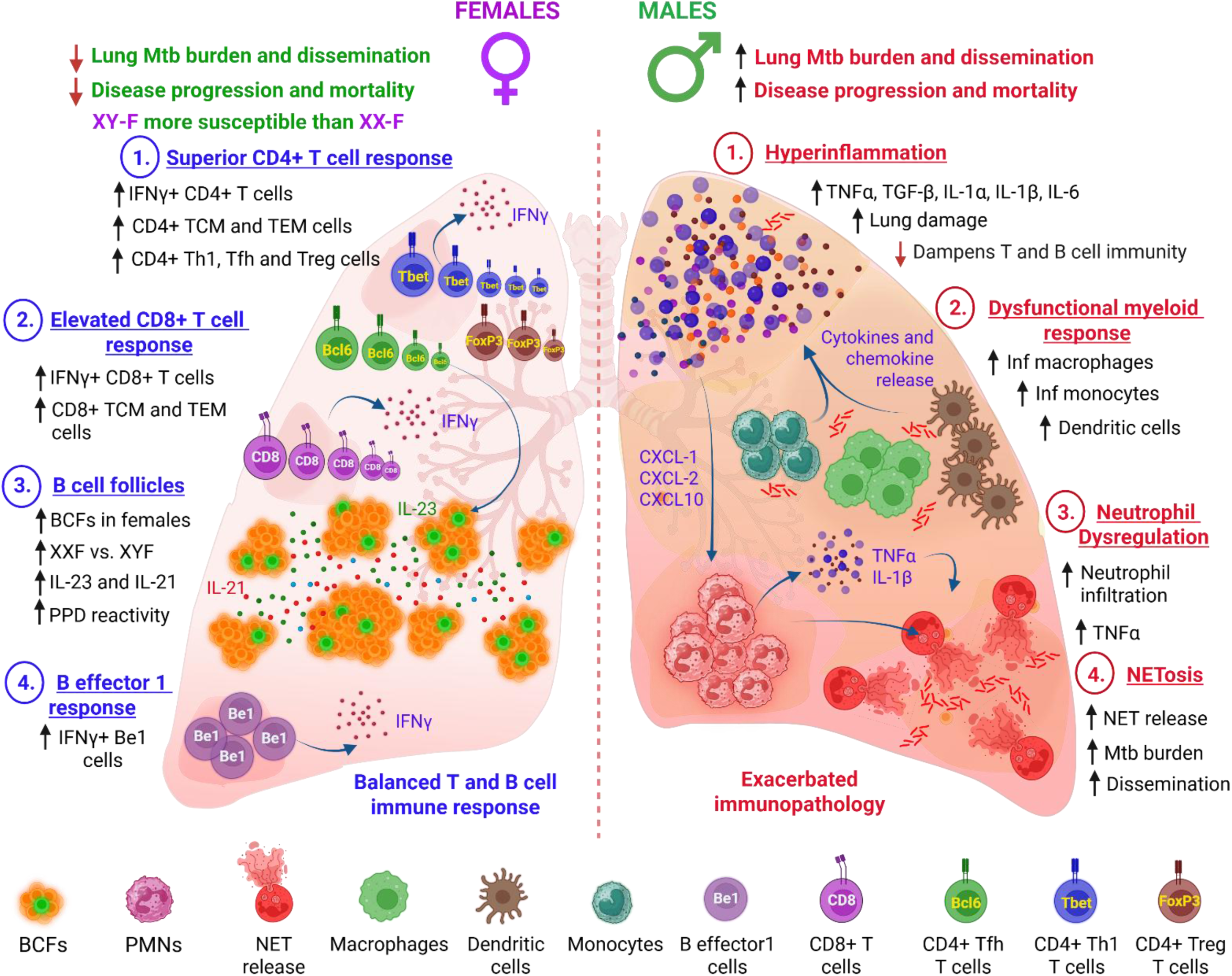
Graphical abstract.

## References

1. World Health Organization (WHO), “Global tuberculosis report 2025” (WHO, 2025).

2. Borgdorff MW, Nagelkerke N, Dye C, Nunn P. Gender and tuberculosis: a comparison of prevalence surveys with notification data to explore sex differences in case detection. The International Journal of Tuberculosis and Lung Disease 4, 123–132 (2000).

3. Fox GJ, Orlova M, Schurr E. Tuberculosis in newborns: the lessons of the “Lübeck Disaster”(1929–1933). PLoS pathogens 12, e1005271 (2016).

4. Hamilton JB, Mestler GE. Mortality and survival: comparison of eunuchs with intact men and women in a mentally retarded population. Journal of Gerontology 24, 395–411 (1969).

5. Dibbern J, Eggers L, Schneider BE. Sex differences in the C57BL/6 model of *Mycobacterium tuberculosis* infection. Scientific reports 7, 10957 (2017).

6. Bini EI, et al. The influence of sex steroid hormones in the immunopathology of experimental pulmonary tuberculosis. PloS one 9, e93831 (2014).

7. Gupta M, Srikrishna G, Klein SL, Bishai WR. Genetic and hormonal mechanisms underlying sex-specific immune responses in tuberculosis. Trends in immunology 43, 640–656 (2022).

8. Klein SL, Flanagan KL. Sex differences in immune responses. Nature Reviews Immunology 16, 626–638 (2016).

9. Hoffmann JP, Liu JA, Seddu K, Klein SL. Sex hormone signaling and regulation of immune function. Immunity 56, 2472–2491 (2023).

10. Moulton VR. Sex hormones in acquired immunity and autoimmune disease. Frontiers in immunology 9, 2279 (2018).

11. Fish EN. The X-files in immunity: sex-based differences predispose immune responses. Nature Reviews Immunology 8, 737–744 (2008).

12. Flanagan KL, Klein SL. Knowledge gaps and research priorities to understand sex differences in immunity. PLoS biology 24, e3003578 (2026).

13. Tomofuji Y, et al. Quantification of escape from X chromosome inactivation with single-cell omics data reveals heterogeneity across cell types and tissues. Cell Genomics 4, (2024).

14. Arnold AP. Four Core Genotypes and XY* mouse models: Update on impact on SABV research. Neuroscience & Biobehavioral Reviews 119, 1–8 (2020).

15. Arnold AP. Conceptual frameworks and mouse models for studying sex differences in physiology and disease: why compensation changes the game. Experimental neurology 259, 2–9 (2014).

16. Hertz D, Dibbern J, Eggers L, von Borstel L, Schneider BE. Increased male susceptibility to *Mycobacterium tuberculosis* infection is associated with smaller B cell follicles in the lungs. Scientific reports 10, 5142 (2020).

17. Swanson RV, et al. Antigen-specific B cells direct T follicular-like helper cells into lymphoid follicles to mediate *Mycobacterium tuberculosis* control. Nature immunology 24, 855–868 (2023).

18. Chowdhury CS, et al. Type I IFN-mediated NET release promotes *Mycobacterium tuberculosis* replication and is associated with granuloma caseation. Cell host & microbe 32, 2092–2111. e2097 (2024).

19. Muefong CN, Sutherland JS. Neutrophils in tuberculosis-associated inflammation and lung pathology. Frontiers in immunology 11, 962 (2020).

20. Öz HH, et al. Recruited monocytes/macrophages drive pulmonary neutrophilic inflammation and irreversible lung tissue remodeling in cystic fibrosis. Cell reports 41, (2022).

21. Chandra P, Grigsby SJ, Philips JA. Immune evasion and provocation by *Mycobacterium tuberculosis*. Nature Reviews Microbiology 20, 750–766 (2022).

22. Harris DP, Goodrich S, Gerth AJ, Peng SL, Lund FE. Regulation of IFN-γ production by B effector 1 cells: essential roles for T-bet and the IFN-γ receptor. The journal of immunology 174, 6781–6790 (2005).

23. Stone SL, et al. T-bet transcription factor promotes antibody-secreting cell differentiation by limiting the inflammatory effects of IFN-γ on B cells. Immunity 50, 1172–1187. e1177 (2019).

24. Bao Y, et al. Identification of IFN-γ-producing innate B cells. Cell research 24, 161–176 (2014).

25. Higgins DM, Sanchez-Campillo J, Rosas-Taraco AG, Lee EJ, Orme IM, Gonzalez-Juarrero M. Lack of IL-10 alters inflammatory and immune responses during pulmonary *Mycobacterium tuberculosis* infection. Tuberculosis 89, 149–157 (2009).

26. Goodman WA, Bedoyan SM, Havran HL, Richardson B, Cameron MJ, Pizarro TT. Impaired estrogen signaling underlies regulatory T cell loss-of-function in the chronically inflamed intestine. Proceedings of the National Academy of Sciences 117, 17166–17176 (2020).

27. Urdahl K, Shafiani S, Ernst J. Initiation and regulation of T-cell responses in tuberculosis. Mucosal immunology 4, 288–293 (2011).

28. Graf R, et al. BCR-dependent lineage plasticity in mature B cells. Science 363, 748–753 (2019).

29. Yang L, et al. Interferon-γ-producing B cells induce the formation of gastric lymphoid follicles after *Helicobacter suis* infection. Mucosal Immunology 8, 279–295 (2015).

30. Krause R, et al. B cell heterogeneity in human tuberculosis highlights compartment-specific phenotype and functional roles. Communications Biology 7, 584 (2024).

31. Wang Q, Nag D, Baldwin SL, Coler RN, McNamara RP. Antibodies as key mediators of protection against *Mycobacterium tuberculosis*. Frontiers in Immunology 15, 1430955 (2024).

32. Bosio CM, Gardner D, Elkins KL. Infection of B cell-deficient mice with CDC 1551, a clinical isolate of *Mycobacterium tuberculosis*: delay in dissemination and development of lung pathology. The Journal of Immunology 164, 6417–6425 (2000).

33. Kozakiewicz L, et al. B cells regulate neutrophilia during *Mycobacterium tuberculosis* infection and BCG vaccination by modulating the interleukin-17 response. PLoS pathogens 9, e1003472 (2013).

34. Khan AR, Hams E, Floudas A, Sparwasser T, Weaver CT, Fallon PG. PD-L1hi B cells are critical regulators of humoral immunity. Nature communications 6, 5997 (2015).

35. Mintz MA, et al. The HVEM-BTLA axis restrains T cell help to germinal center B cells and functions as a cell-extrinsic suppressor in lymphomagenesis. Immunity 51, 310–323. e317 (2019).

36. Khader SA, et al. IL-23 is required for long-term control of *Mycobacterium tuberculosis* and B cell follicle formation in the infected lung. The Journal of Immunology 187, 5402–5407 (2011).

37. Majdic G, Tobet S. Cooperation of sex chromosomal genes and endocrine influences for hypothalamic sexual differentiation. Frontiers in Neuroendocrinology 32, 137–145 (2011).

38. Arnold AP. Sexual differentiation of brain and other tissues: Five questions for the next 50 years. Hormones and behavior 120, 104691 (2020).

39. Robinson DP, Klein SL. Pregnancy and pregnancy-associated hormones alter immune responses and disease pathogenesis. Hormones and behavior 62, 263–271 (2012).

40. Hall OJ, Klein SL. Progesterone-based compounds affect immune responses and susceptibility to infections at diverse mucosal sites. Mucosal Immunology 10, 1097–1107 (2017).

41. Khan D, Ansar Ahmed S. The immune system is a natural target for estrogen action: opposing effects of estrogen in two prototypical autoimmune diseases. Frontiers in immunology 6, 635 (2016).

42. Goldman AL, Bhasin S, Wu FC, Krishna M, Matsumoto AM, Jasuja R. A reappraisal of testosterone’s binding in circulation: physiological and clinical implications. Endocrine reviews 38, 302–324 (2017).

43. Panten J, et al. Four Core Genotypes mice harbour a 3.2 MB XY translocation that perturbs Tlr7 dosage. Nature Communications 15, 8814 (2024).

44. Burgoyne PS, Arnold AP. A primer on the use of mouse models for identifying direct sex chromosome effects that cause sex differences in non-gonadal tissues. Biology of sex differences 7, 68 (2016).

45. Bolger AM, Lohse M, Usadel B. Trimmomatic: a flexible trimmer for Illumina sequence data. Bioinformatics 30, 2114–2120 (2014).

46. Dobin A, et al. STAR: ultrafast universal RNA-seq aligner. Bioinformatics 29, 15–21 (2013).

47. Love MI, Huber W, Anders S. Moderated estimation of fold change and dispersion for RNA-seq data with DESeq2. Genome biology 15, 550 (2014).

