## Supplementary Figures for "Gonadal regulation of sex-specific immunity in tuberculosis: enhanced lymphocyte function in females and dysfunctional myeloid responses in males"

**FigS1.**

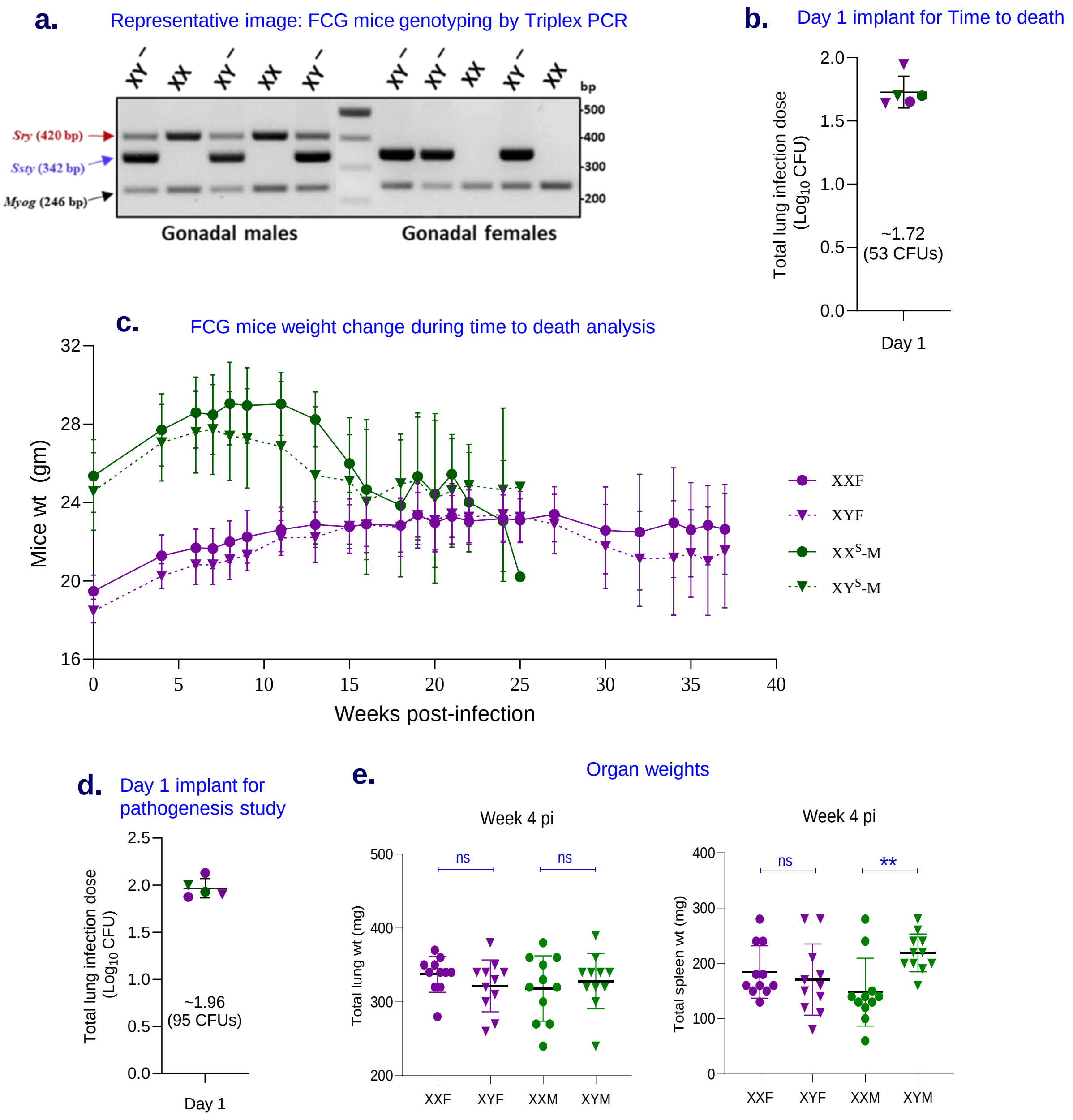


**Figure S1. FCG mice genotyping and Mtb infection studies. (a).** Crossing XXF with $XY^{-Sry}$M produces the four core genotypes- XXF and XYF with ovaries, and XXM and XYM with testes. A 2% Agarose gel electrophoresis depicting FCG genotyping by triplex PCR, for amplifying - *Ssty* = Y chromosome; *Sry* = *Sry*+ Chr3; *Myog* = loading control. **(b).** FCG mice were infected with Mtb H37Rv at 8–10 weeks of age by aerosol route. Achieved day 1 lung Mtb burden for time-to-death study (n=5). **(c).** FCG mice body weight change kinetics during the time to death analysis, after Mtb infection (at day 0, n=15). **(d).** Day 1 implant of FCG mice, during pathogeneisis study. **(e).** Gross lung and spleen weights of FCG mice (n = 11) at 4 wpi. **(f).** Serum estradiol (n=4 for week 0, and n=12 for week 4 and 12, post-infection) and testosterone levels (n=9 for week 0, and n=12 for week 4 and 12, post-infection). **(g)**. Survival kinetics of Mtb H37Rv in IFNγ-activated BMDMs isolated from FCG mice (MOI:1; n=3/group). These BMDMs were lysed at indicated time intervals and plated on 7H11 selection plates for CFU enumeration. **(h).** Normalized gene expression measured by qPCR at 6 h post Mtb infection (MOI:5). Data shown as fold change relative to uninfected groups, using the $2^{-\Delta\Delta Ct}$ method (n = 6). Data were plotted as mean ± SEM. Statistical significance was calculated using two-way ANOVA. *$P < 0.05$, ** $P < 0.01$, *** $P < 0.001$, **** $P < 0.0001$ and ns for no significance.

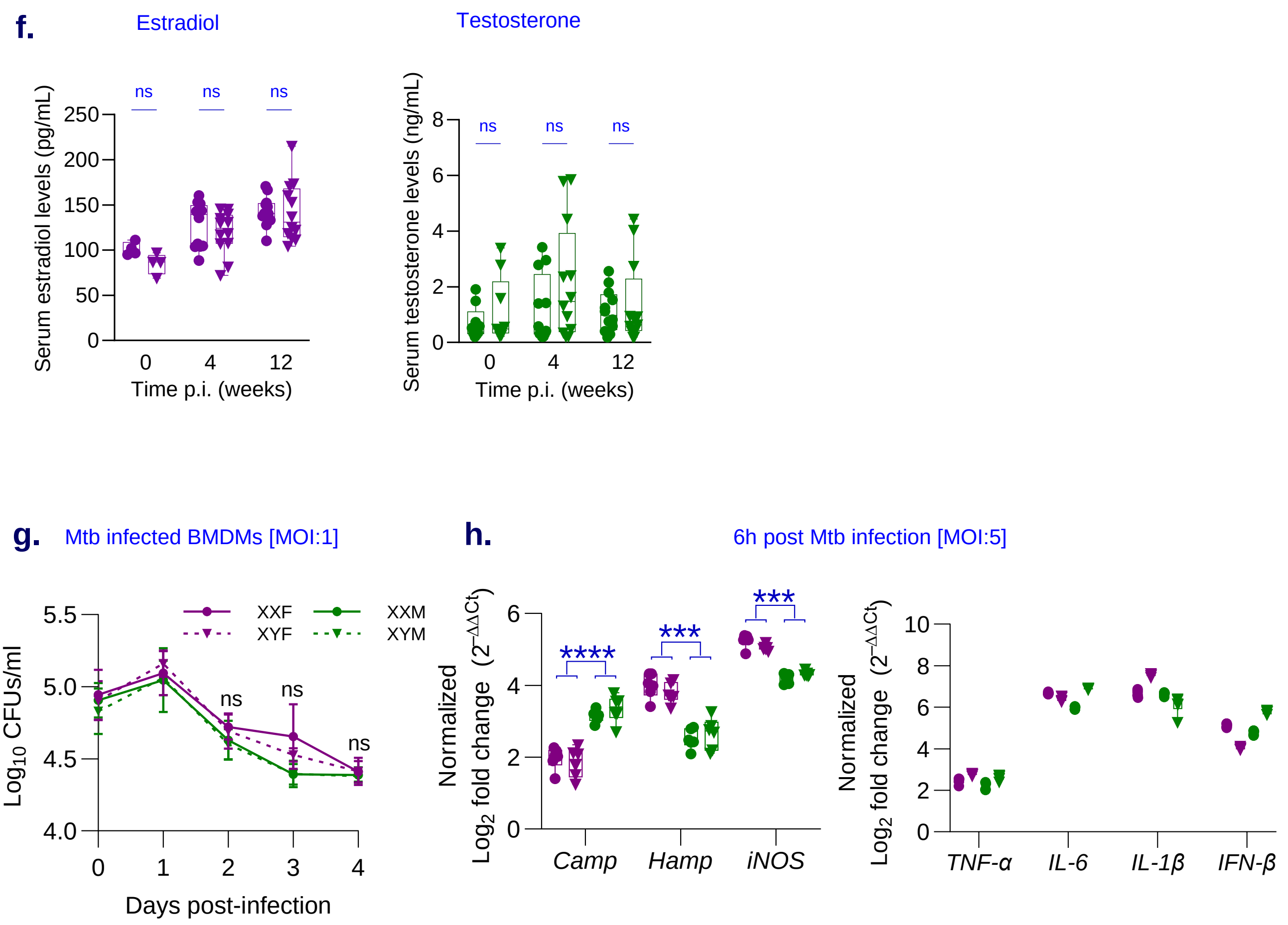
f.
Estradiol
Serum estradiol levels (pg/mL)
Time p.i. (weeks)
ns
Testosterone
Serum testosterone levels (ng/mL)
Time p.i. (weeks)
ns
g.
Mtb infected BMDMs [MOI:1]
XXF
XYF
XXM
XYM
Log10 CFUs/ml
Days post-infection
ns
h.
6h post Mtb infection [MOI:5]
Normalized
Log2 fold change (2−ΔΔCt)
Camp
Hamp
iNOS
****
***
TNF-α
IL-6
IL-1β
IFN-β

**FigS2.**

**a.**

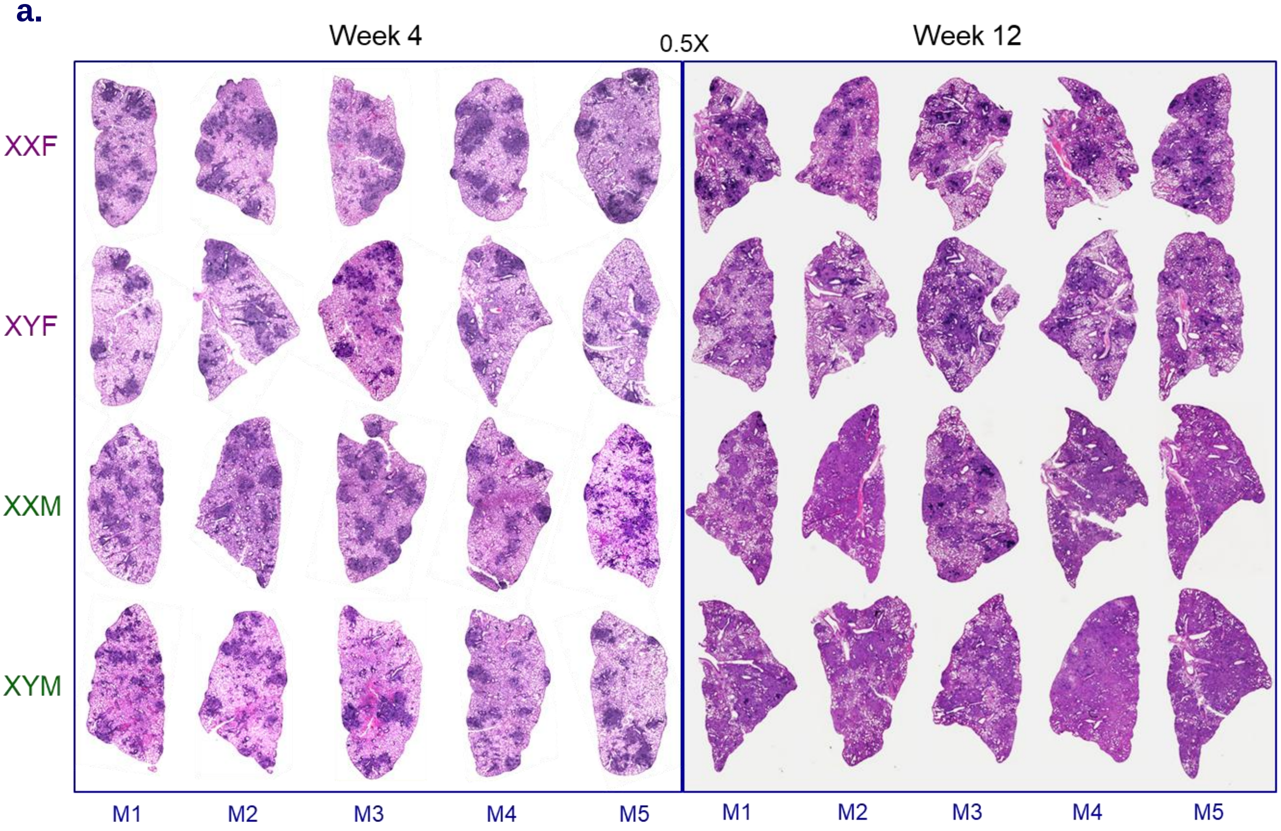


**Figure S2. FCG mice lung histopathology and cytokine/chemokine profiling at week 4 and 12 weeks post-infection with Mtb H37Rv. (a).** Lung histopathology of Mtb-infected FCG mice was performed at 4 and 12 wpi. The lungs were formalin-fixed, sectioned, and stained with hematoxylin and eosin (H&E). Five representative images/group. Lung inflammatory responses in gonadal males and females were measured by quantifying cytokine and chemokine levels in lung lysates after 4 and 12 wpi post infection. **(b-m).** Bar graph visualization of kinetics of pro-inflammatory and anti-inflammatory cytokines. **(n-ac).** Bar graph visualization of kinetics of various chemokines analyzed in this study (n=6). Data were plotted as mean ± SEM. Statistical significance was calculated using two-way ANOVA. *$P < 0.05$, ** $P < 0.01$, *** $P < 0.001$ and ns for no significance.

**FigS2.**

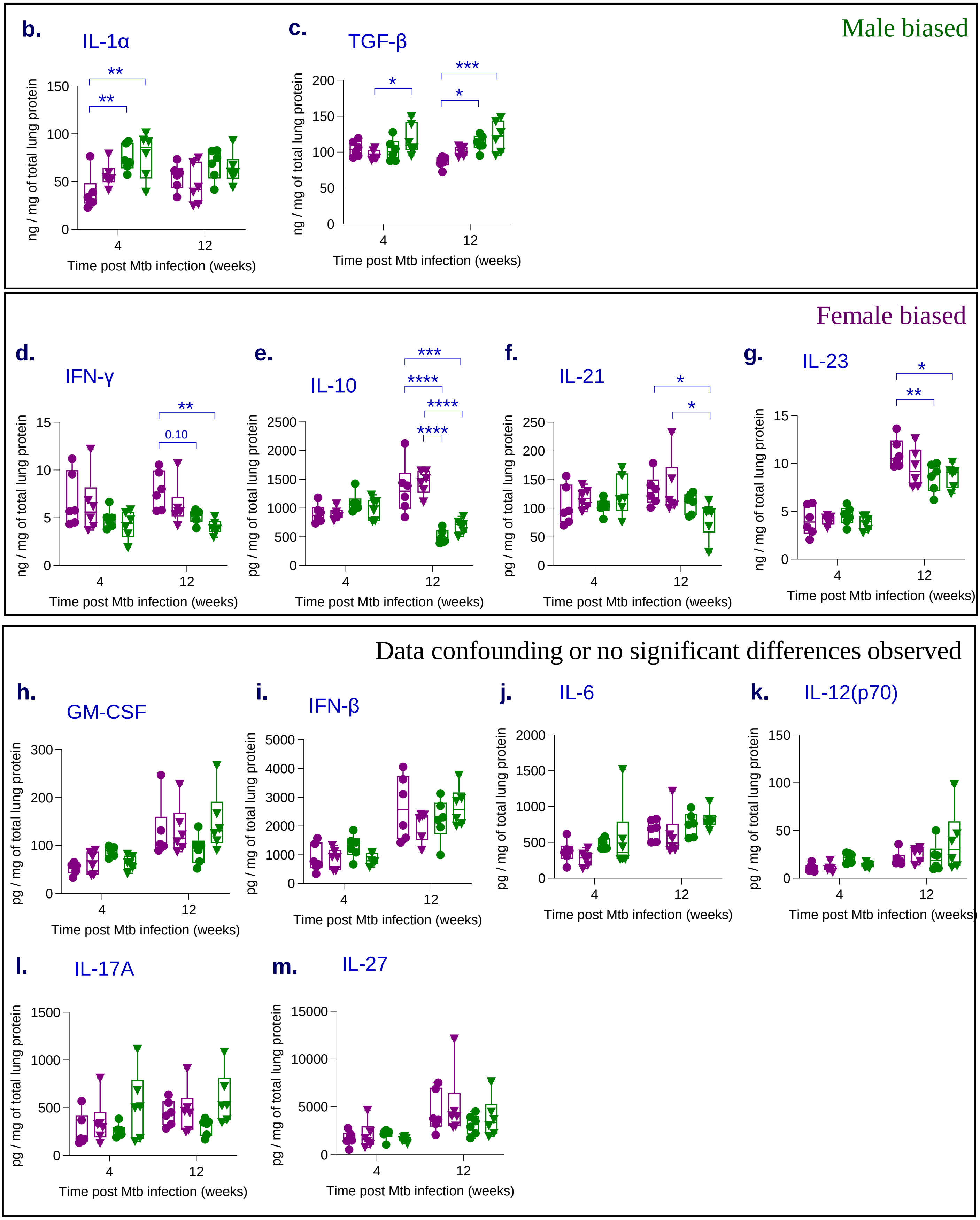

Male biased
b.
IL-1α
ng / mg of total lung protein
Time post Mtb infection (weeks)
c.
TGF-β
ng / mg of total lung protein
Time post Mtb infection (weeks)
Female biased
d.
IFN-γ
0.10
ng / mg of total lung protein
Time post Mtb infection (weeks)
e.
IL-10
pg / mg of total lung protein
Time post Mtb infection (weeks)
f.
IL-21
pg / mg of total lung protein
Time post Mtb infection (weeks)
g.
IL-23
ng / mg of total lung protein
Time post Mtb infection (weeks)
Data confounding or no significant differences observed
h.
GM-CSF
pg / mg of total lung protein
Time post Mtb infection (weeks)
i.
IFN-β
pg / mg of total lung protein
Time post Mtb infection (weeks)
j.
IL-6
pg / mg of total lung protein
Time post Mtb infection (weeks)
k.
IL-12(p70)
pg / mg of total lung protein
Time post Mtb infection (weeks)
l.
IL-17A
pg / mg of total lung protein
Time post Mtb infection (weeks)
m.
IL-27
pg / mg of total lung protein
Time post Mtb infection (weeks)

**FigS2.**

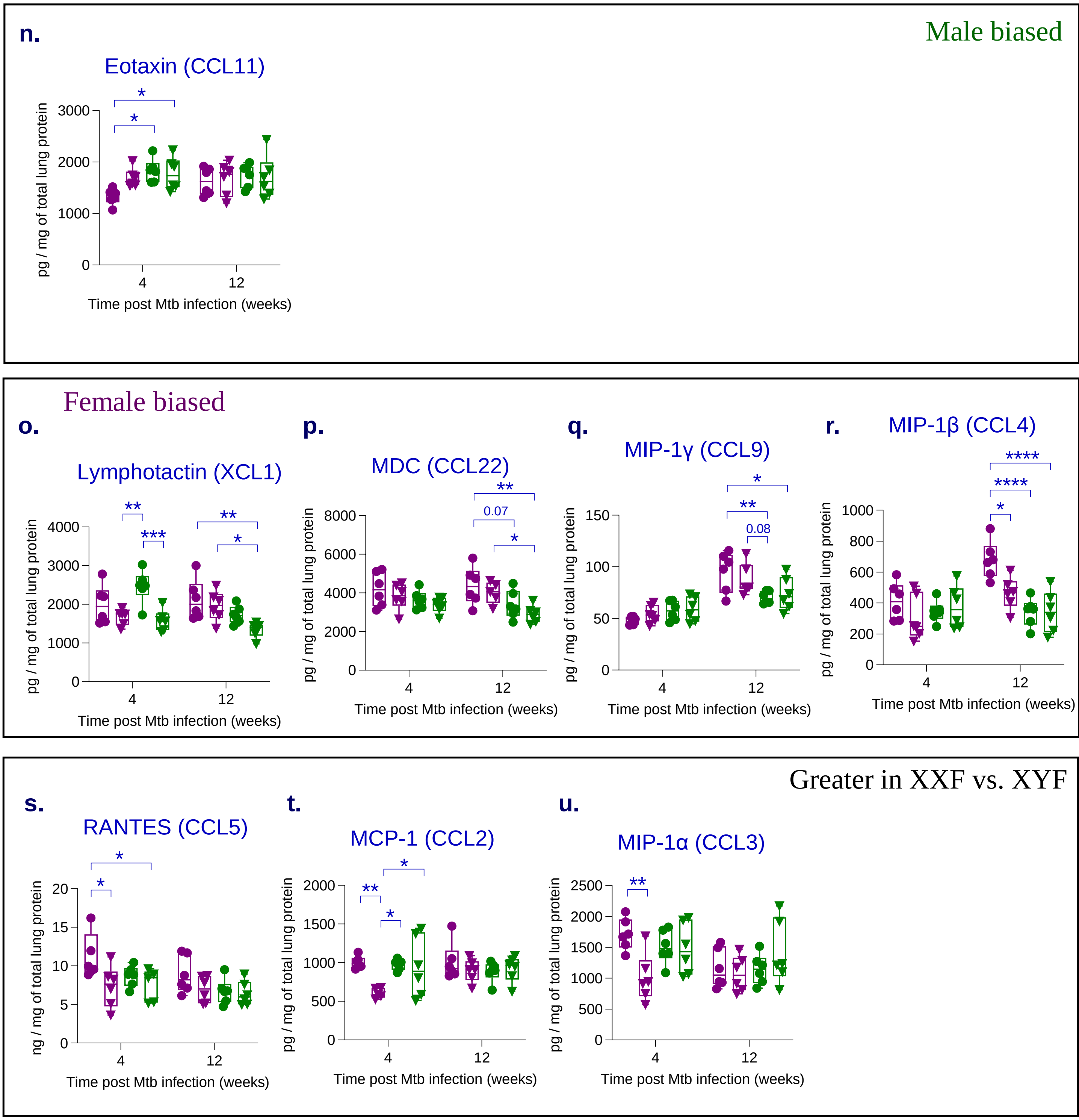

Male biased
n.
Eotaxin (CCL11)
pg / mg of total lung protein
Time post Mtb infection (weeks)
Female biased
o.
Lymphotactin (XCL1)
p.
MDC (CCL22)
q.
MIP-1γ (CCL9)
r.
MIP-1β (CCL4)
0.07
0.08
Greater in XXF vs. XYF
s.
RANTES (CCL5)
ng / mg of total lung protein
t.
MCP-1 (CCL2)
u.
MIP-1α (CCL3)

No significant differences observed

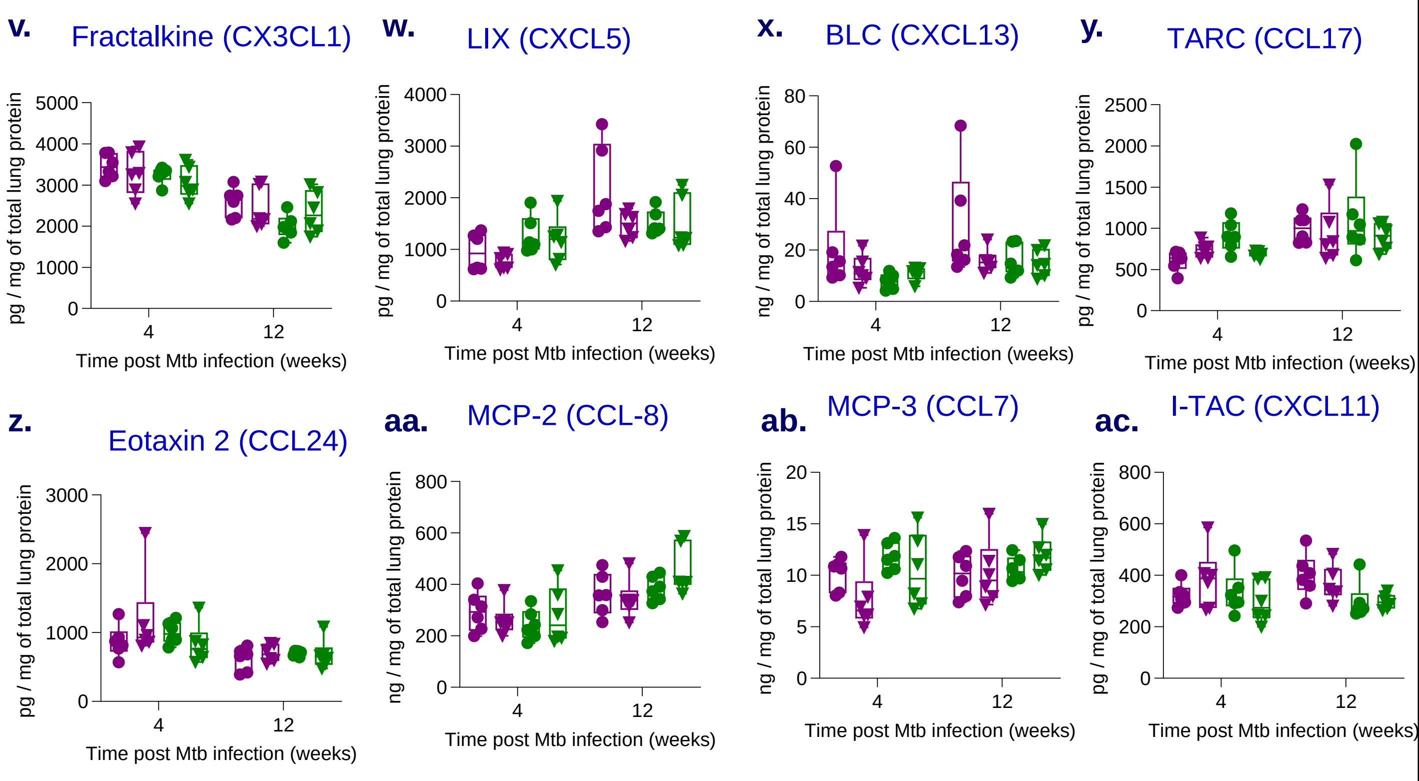

**FigS3.**

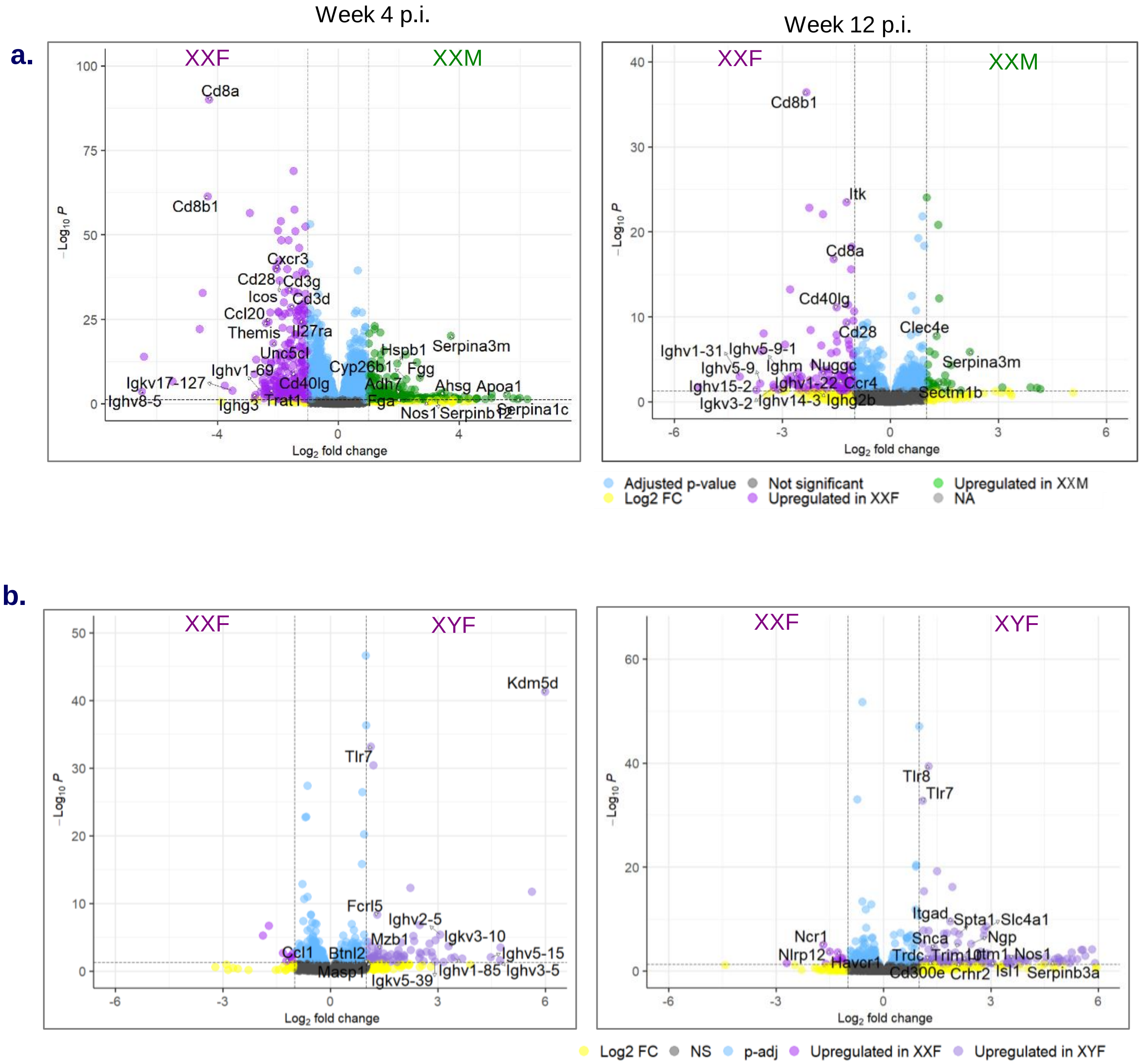

Week 4 p.i.
Week 12 p.i.
a.
XXF
XXM
Cd8a
Cd8b1
Cxcr3
Cd28
Cd3g
Icos
Cd3d
Ccl20
Themis
Il27ra
Unc5cl
Hspb1
Serpina3m
Cyp26b1
Fgg
Ighv1-69
Igkv17-127
Cd40lg
Adh7
Ahsg
Apoa1
Ighv8-5
Ighg3
Trat1
Fga
Nos1
Serpinb12
Serpina1c
Log2 fold change
−Log10 P
Itk
Cd40lg
Clec4e
Ighv1-31
Ighv5-9-1
Ighv5-9
Ighm
Nuggc
Ighv15-2
Ighv1-22
Ccr4
Igkv3-2
Ighv14-3
Ighg2b
Sectm1b
Adjusted p-value
Log2 FC
Not significant
Upregulated in XXF
Upregulated in XXM
NA
b.
XYF
Kdm5d
Tlr7
Fcrl5
Ighv2-5
Mzb1
Igkv3-10
Ccl1
Btnl2
Ighv5-15
Masp1
Ighv1-85
Ighv3-5
Igkv5-39
Tlr8
Itgad
Spta1
Slc4a1
Ncr1
Snca
Ngp
Nlrp12
Trdc
Trim10
Itm1
Nos1
Havcr1
Cd300e
Crhr2
Isl1
Serpinb3a
NS
p-adj
Upregulated in XYF

c.

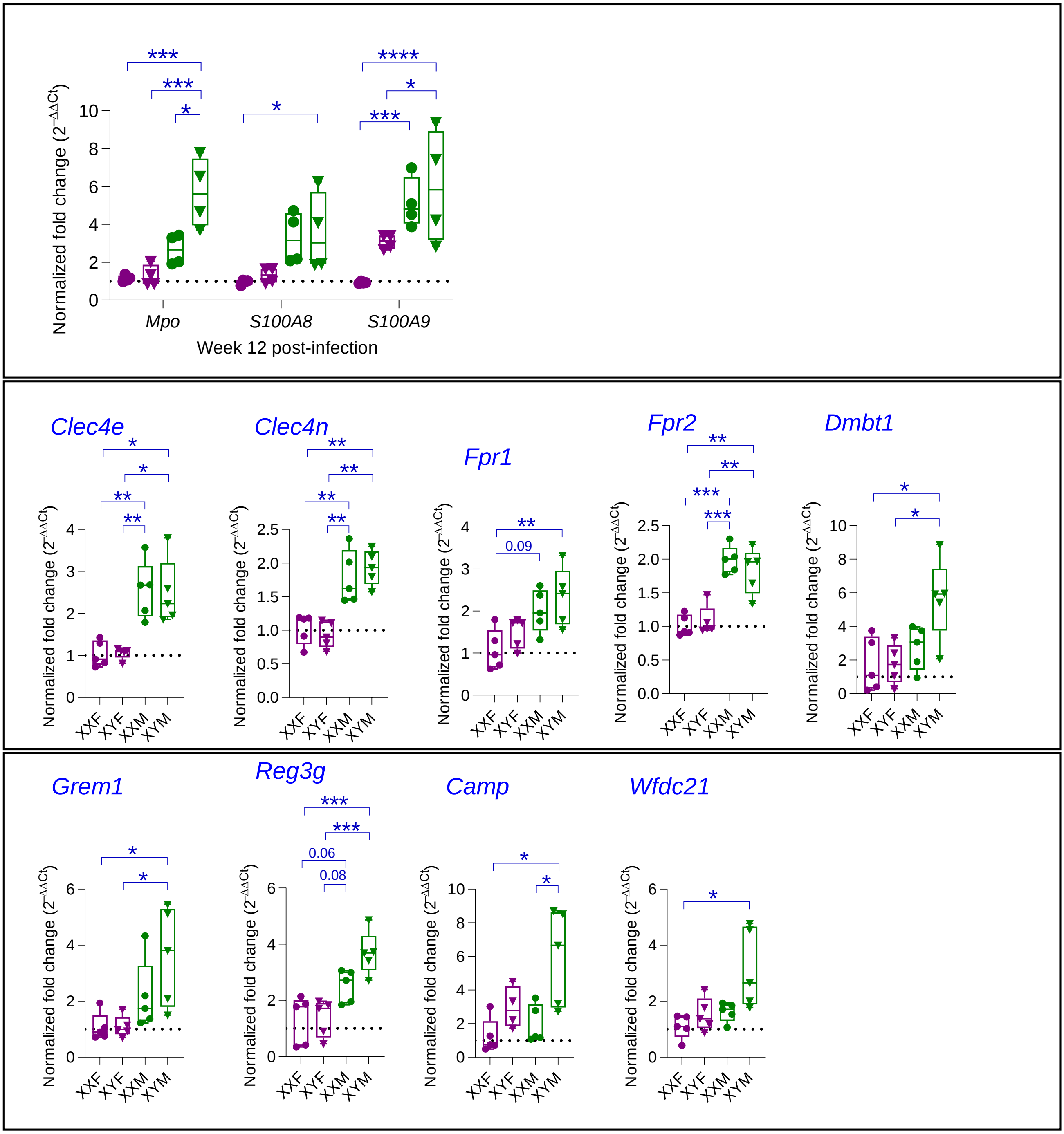

Normalized fold change ($2^{-\Delta\Delta Ct}$)
Mpo
S100A8
S100A9
Week 12 post-infection
***
***
*
*
****
*
***
Clec4e
Clec4n
Fpr1
Fpr2
Dmbt1
0.09
Grem1
Reg3g
Camp
Wfdc21
0.06
0.08
XXF
XYF
XXM
XYM

d.

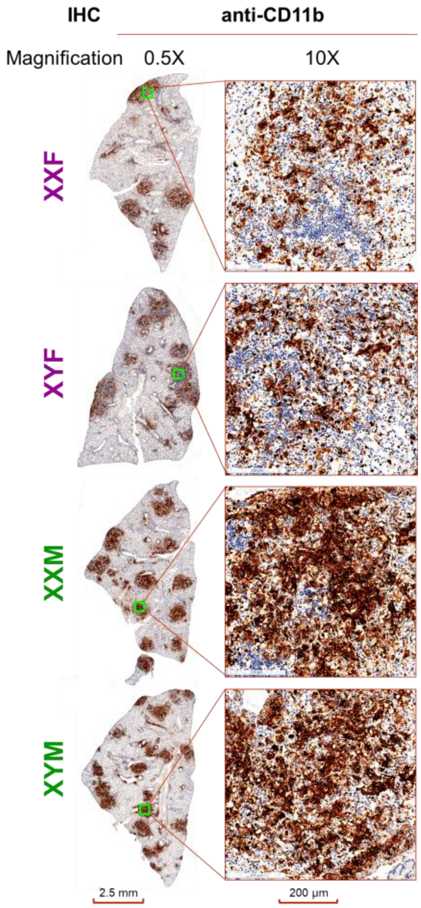


IHC staining with anti-CD11b and anti-F4/80 antibodies of Mtb-infected FCG mice lungs (4X magnification), week 4 post-infection

e.

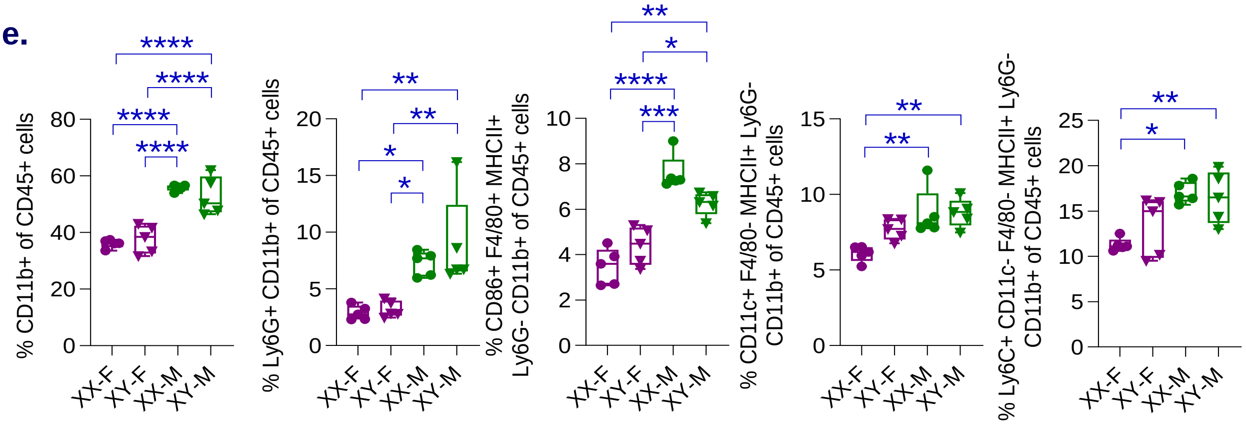

f.

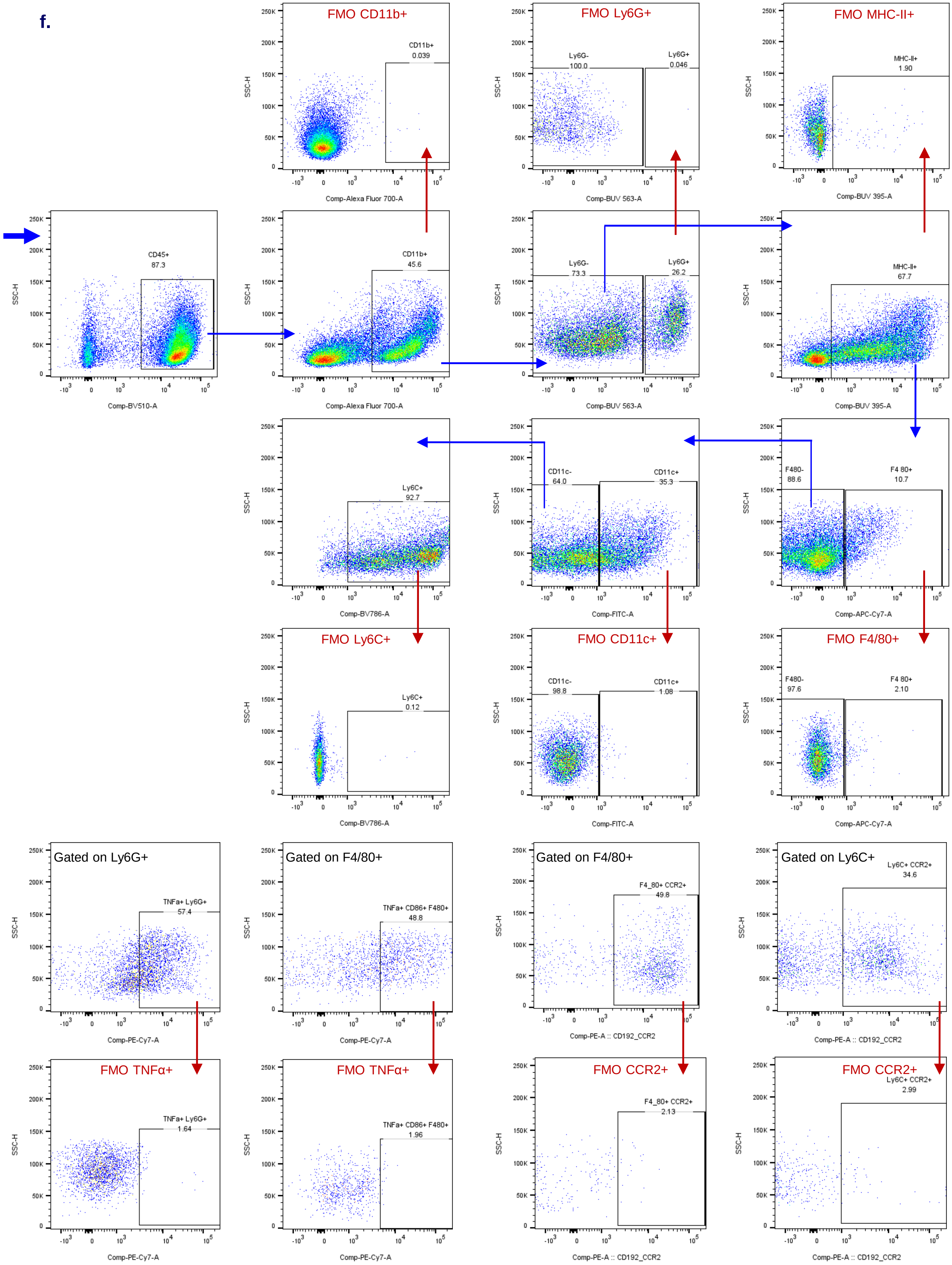

FMO CD11b+
CD11b+
0.039
Comp-Alexa Fluor 700-A
FMO Ly6G+
Ly6G-
100.0
Ly6G+
0.046
Comp-BUV 563-A
FMO MHC-II+
MHC-II+
1.90
Comp-BUV 395-A
CD45+
87.3
Comp-BV510-A
CD11b+
45.6
Comp-Alexa Fluor 700-A
Ly6G-
73.3
Ly6G+
26.2
Comp-BUV 563-A
MHC-II+
67.7
Comp-BUV 395-A
Ly6C+
92.7
Comp-BV786-A
CD11c-
64.0
CD11c+
35.3
Comp-FITC-A
F480-
88.6
F4 80+
10.7
Comp-APC-Cy7-A
FMO Ly6C+
Ly6C+
0.12
Comp-BV786-A
FMO CD11c+
CD11c-
98.8
CD11c+
1.08
Comp-FITC-A
FMO F4/80+
F480-
97.6
F4 80+
2.10
Comp-APC-Cy7-A
Gated on Ly6G+
TNFa+ Ly6G+
57.4
Comp-PE-Cy7-A
Gated on F4/80+
TNFa+ CD86+ F480+
48.8
Comp-PE-Cy7-A
Gated on F4/80+
F4_80+ CCR2+
49.8
Comp-PE-A :: CD192_CCR2
Gated on Ly6C+
Ly6C+ CCR2+
34.6
Comp-PE-A :: CD192_CCR2
FMO TNFα+
TNFa+ Ly6G+
1.64
Comp-PE-Cy7-A
FMO TNFα+
TNFa+ CD86+ F480+
1.96
Comp-PE-Cy7-A
FMO CCR2+
F4_80+ CCR2+
2.13
Comp-PE-A :: CD192_CCR2
FMO CCR2+
Ly6C+ CCR2+
2.99
Comp-PE-A :: CD192_CCR2
SSC-H

g.

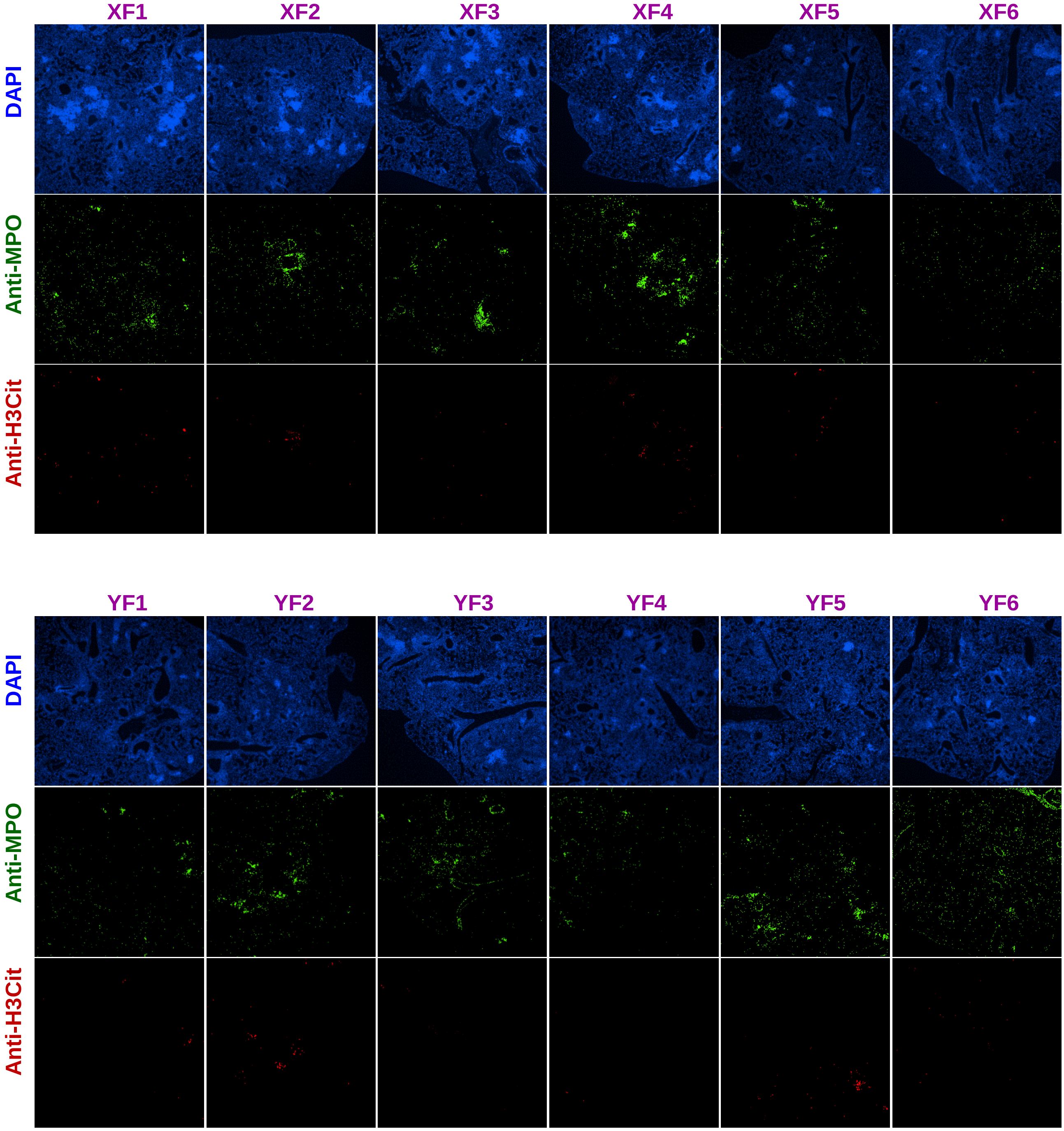
XF1
XF2
XF3
XF4
XF5
XF6
DAPI
Anti-MPO
Anti-H3Cit
YF1
YF2
YF3
YF4
YF5
YF6
DAPI
Anti-MPO
Anti-H3Cit

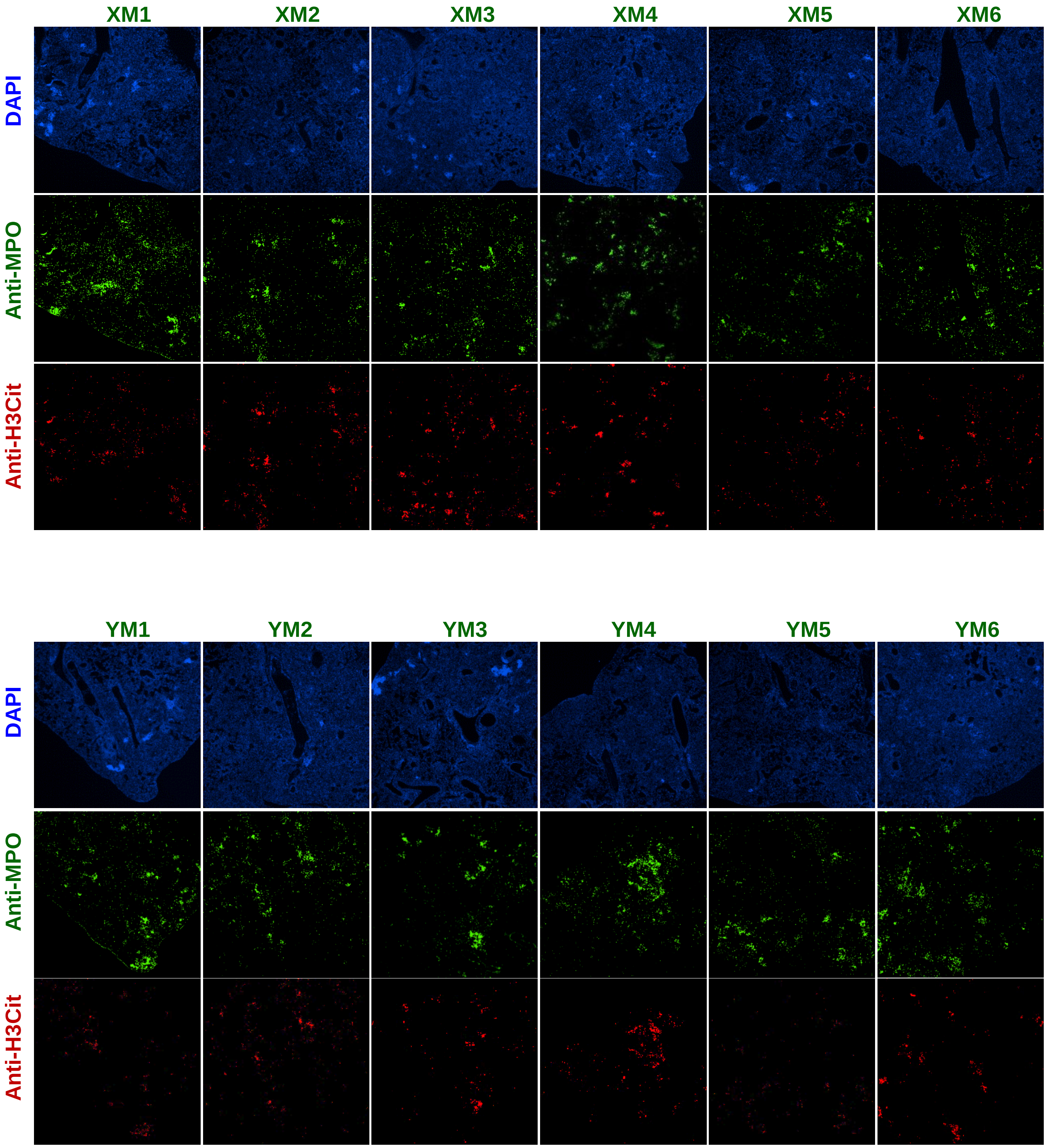
XM1
XM2
XM3
XM4
XM5
XM6
DAPI
Anti-MPO
Anti-H3Cit
YM1
YM2
YM3
YM4
YM5
YM6
DAPI
Anti-MPO
Anti-H3Cit

**h.**

NETosis

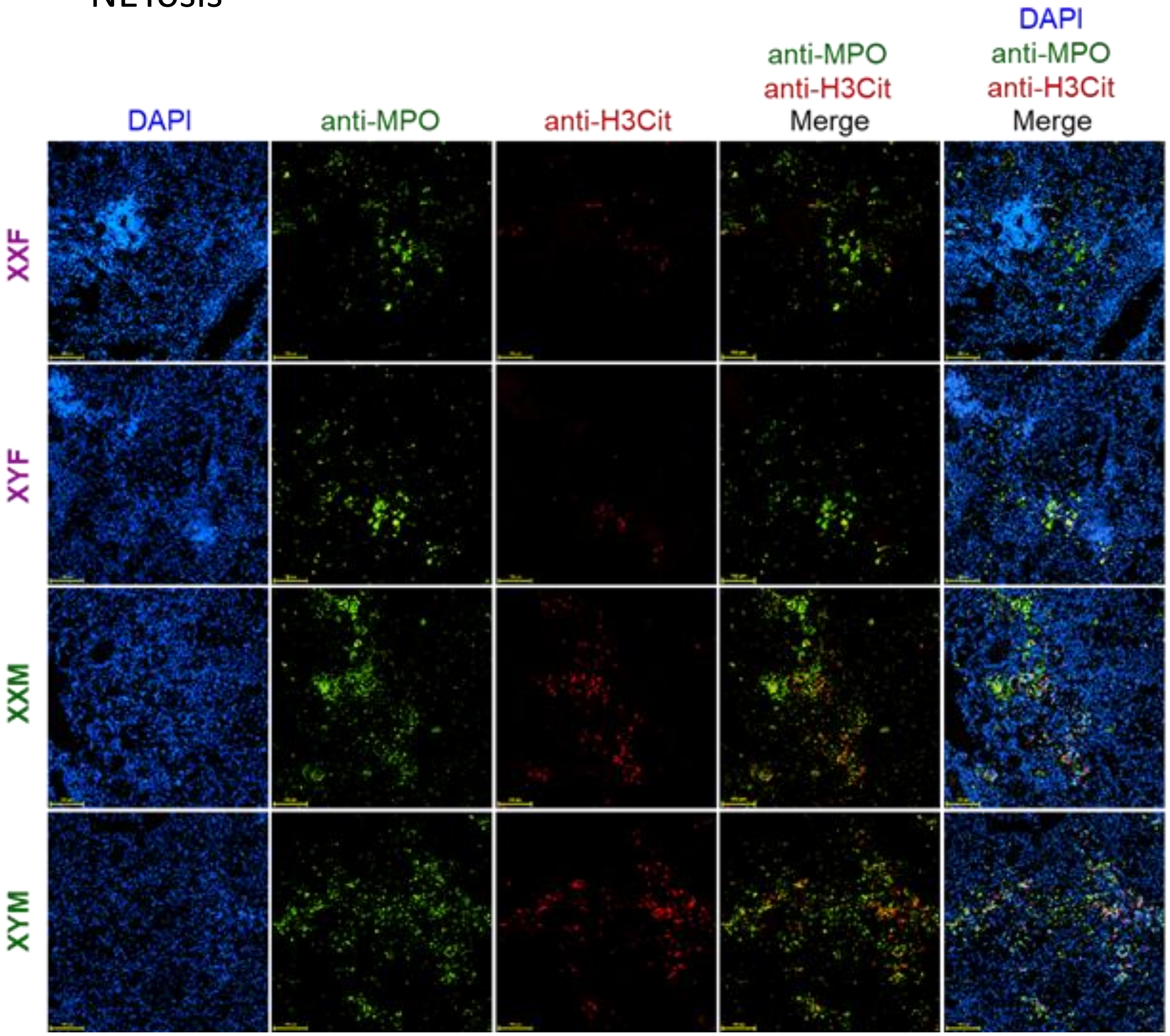
DAPI
anti-MPO
anti-H3Cit
anti-MPO
anti-H3Cit
Merge
DAPI
anti-MPO
anti-H3Cit
Merge
XXF
XYF
XXM
XYM

**Figure S3. Elevated yet dysfunctional myeloid response in males after Mtb infection.** Volcano plot representing differential gene expression in – **(a).** XXF vs. XXM and **(b).** XXF vs. XYF lungs after 4 and 12 weeks post-Mtb infection (n=5). The negative base-10 logarithm of P-adjusted values is plotted on the Y-axis, and fold change (Log2) is plotted on the X-axis. Purple indicates transcripts up-regulated in XXF, green indicates transcripts up-regulated in XXM, while lavender indicates transcripts up-regulated in XYF, and blue indicates no significantly differentially expressed genes (Log2 FC >1, $P < 0.05$). **(c).** Validation of RNASeq data, showing upregulation of myeloid-associated genes in gonadal males. Normalized gene expression of key genes measured by qPCR at 12 wpi. Data shown as fold change relative to XXF, using $2^{-\Delta\Delta Ct}$ method. For PMN associated genes, viz., *Mpo*, *S100a8* and *S100a9*, n=4. For all the others genes n=5. **(d).** At week 4 post-infection formalin-fixed lung tissue were sectioned, and stained with anti-CD11b antibodies. Representative CD11b-stained lung sections at 0.5x magnification. Insets in sections depict CD11b+ stained areas in localized granulomas at 10x magnification. As shown in Fig 3g-j, single-cell suspensions from FCG mice lungs were stimulated for 4 h (detailed in the Methods), and TNF-α expression on different myeloid subsets was measured by multicolor flow cytometry (n=5). **(e).** Depicts the total cell frequencies of each myeloid subset related to Fig 3g-j. Flow cytometry analysis showing the frequencies of CD11b+ all myeloid cells, PMNs/neutrophils (Ly6G+ CD11b+), M1 macrophages (CD86+ F4/80+ MHC-II+), dendritic cells (CD11c+ F4/80- MHC-II+), and monocytes (Ly6C+ F4/80- CD11c- MHC-II+), at week 4 post infection with Mtb. No data for 12 wpi, due to loss of samples during processing (n = 5). **(f).** Two-dimensional gating strategies for identification of different myeloid cell subsets by flow cytometry. All flow cytometry antibodies were titrated to determine the optimal concentration that provided maximal specificity with minimal spectral spillover. The gating strategy was established using single-stained controls and fluorescence-minus-one (FMO) controls. Gates are depicted in blue/black and respective FMO controls are labeled in red. **(g).** Immunofluorescence staining of Mtb-infected FCG mouse lungs at 12 wpi (same sections as used in Fig. 2a), stained with fluorescently labeled antibodies to detect myeloperoxidase (MPO in green, for probing neutrophilic granules), citrullinated histone H3 (H3Cit labelling, in red), and nuclear counterstain (DAPI to stain DNA, in blue). n=6/group. **(h).** Same as Fig. 3n. Representative 20X scans, but also showing the merged image with DAPI, anti-MPO and anti-H3Cit sections. Each dot represents an individual mouse. Data are represented as mean ± SEM values. Statistical significance was calculated using two-way ANOVA. $^*P < 0.05$, $^{**} P < 0.01$, $^{***} P < 0.001$ and ns for no significance.

**FigS4.**

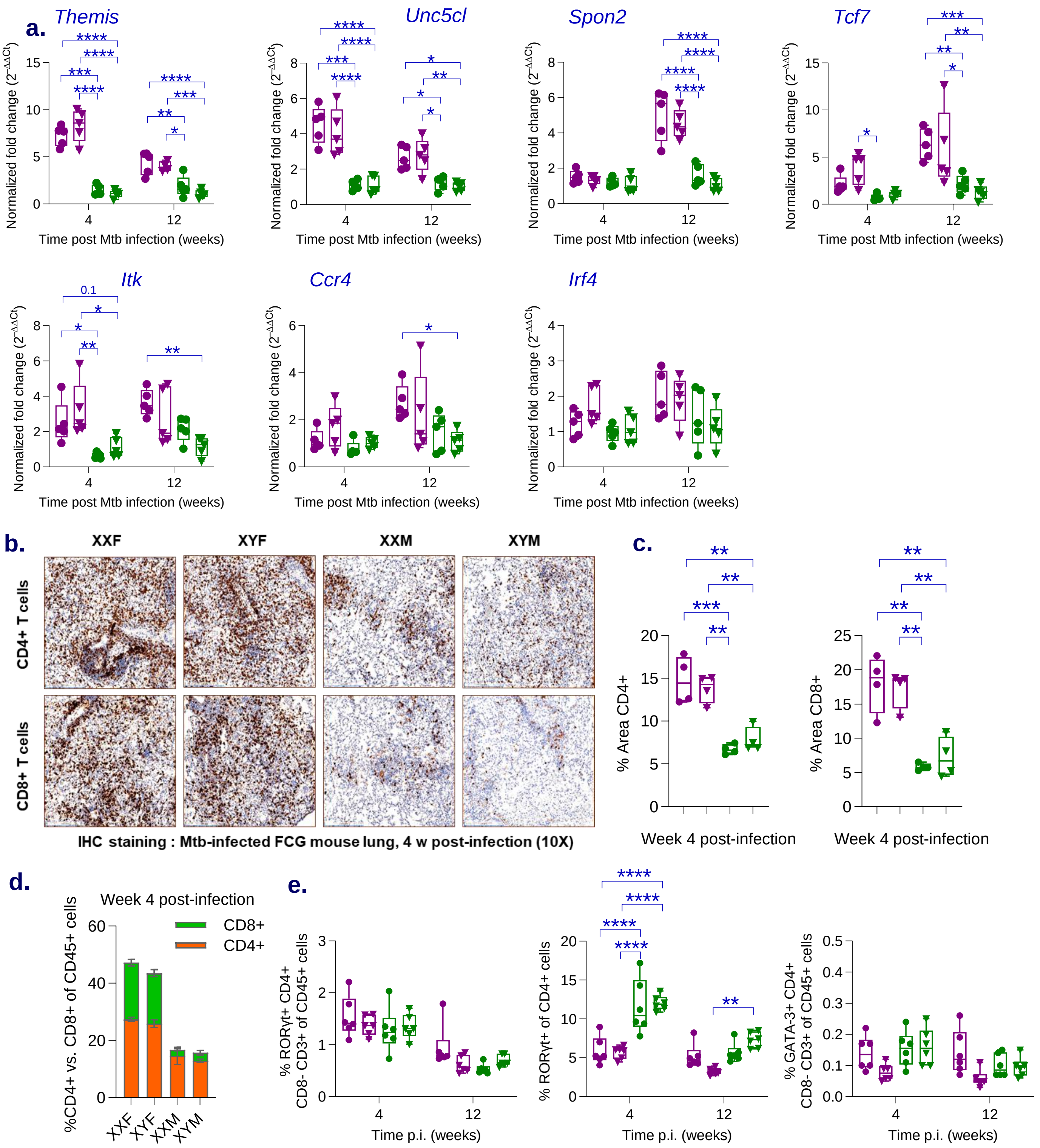

a.
Themis
Unc5cl
Spon2
Tcf7
Itk
Ccr4
Irf4
Normalized fold change (2^−ΔΔCt)
Time post Mtb infection (weeks)
0.1
b.
XXF
XYF
XXM
XYM
CD4+ T cells
CD8+ T cells
IHC staining : Mtb-infected FCG mouse lung, 4 w post-infection (10X)
c.
% Area CD4+
% Area CD8+
Week 4 post-infection
d.
Week 4 post-infection
CD8+
CD4+
%CD4+ vs. CD8+ of CD45+ cells
XXF
XYF
XXM
XYM
e.
% RORγt+ CD4+ CD8- CD3+ of CD45+ cells
% RORγt+ of CD4+ cells
% GATA-3+ CD4+ CD8- CD3+ of CD45+ cells
Time p.i. (weeks)

**Figure S4. Gonadal females mount a superior T cell response to Mtb infection. (a).** Validation of RNASeq data, showing upregulation of lymphoid-associated genes in gonadal females. Normalized gene expression of key genes measured by qPCR at 4 and 12 wpi. Data shown as fold change relative to XYM, using $2^{-\Delta\Delta Ct}$ method. For all the genes n=5/group. **(b).** At week 4 post-infection formalin-fixed lung tissue were sectioned, and stained with anti-CD4 and anti-CD8 antibodies. Representative CD4 and CD8-stained lung sections at 10x magnification. **(c).** Quantification of total CD4+ and CD8+ area/lung section at 4 wpi (n=4). **(d).** Distribution of CD4+ and CD8+ T cells in FCG males and females at 4 wpi. **(e).** Lung immunophenotyping showing- frequencies of RORγt+ CD4+ Th17 cells and GATA3+ CD4+ Th2 cells (n=6). Each dot represents an individual mouse. Data are represented as mean ± SEM values. Statistical significance was calculated using two-way ANOVA. *$P < 0.05$, ** $P < 0.01$, *** $P < 0.001$ and ns for no significance.

**FigS5.**

**a.**

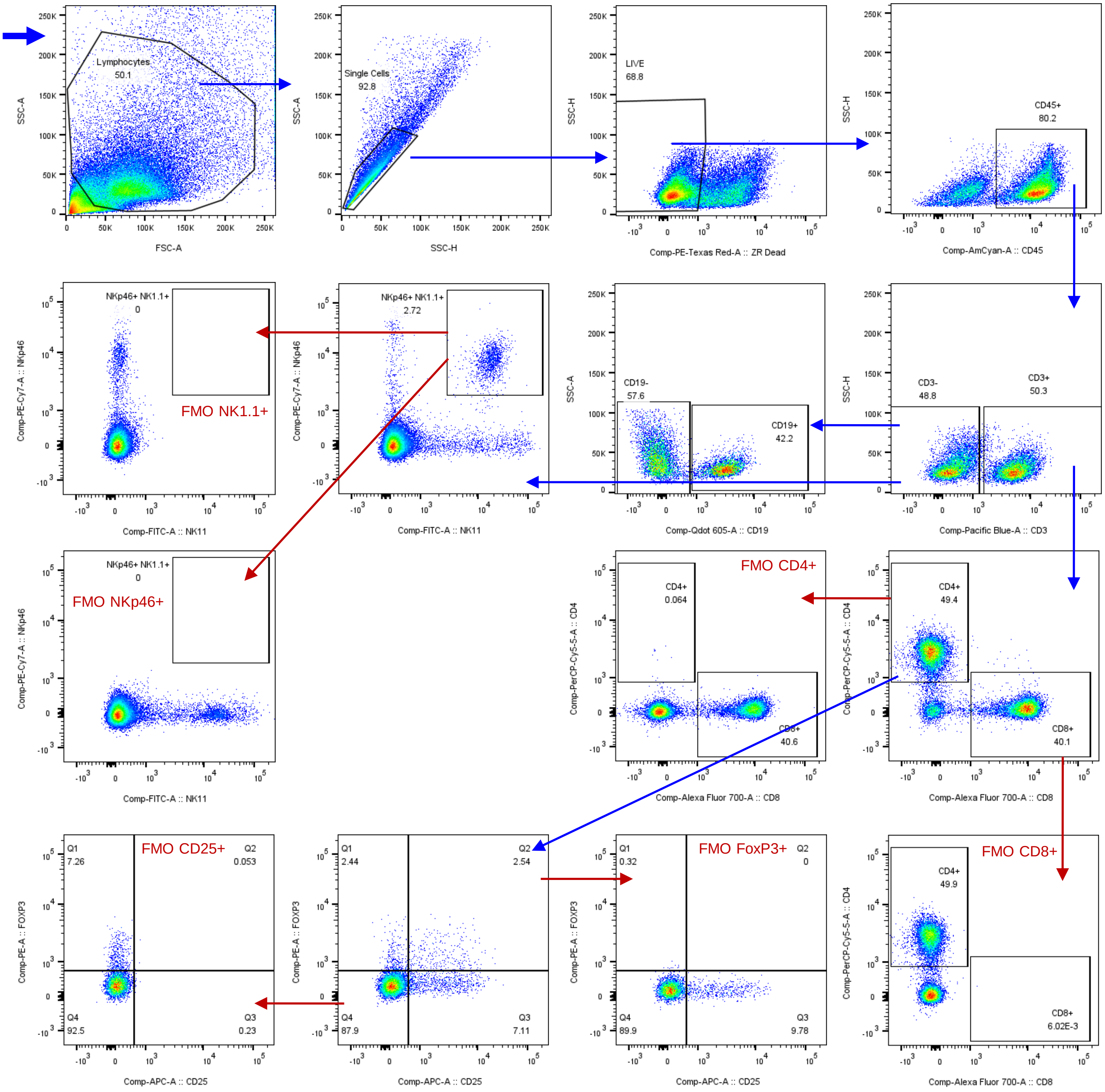
Lymphocytes
50.1
Single Cells
92.8
LIVE
68.8
CD45+
80.2
FMO NK1.1+
NKp46+ NK1.1+
0
NKp46+ NK1.1+
2.72
CD19-
57.6
CD19+
42.2
CD3-
48.8
CD3+
50.3
FMO NKp46+
FMO CD4+
CD4+
0.064
CD8+
40.6
CD4+
49.4
CD8+
40.1
FMO CD25+
Q1 7.26
Q2 0.053
Q4 92.5
Q3 0.23
Q1 2.44
Q2 2.54
Q4 87.9
Q3 7.11
FMO FoxP3+
Q1 0.32
Q2 0
Q4 89.9
Q3 9.78
FMO CD8+
CD4+
49.9
CD8+
6.02E-3
FSC-A
SSC-A
SSC-H
Comp-PE-Texas Red-A :: ZR Dead
Comp-AmCyan-A :: CD45
Comp-PE-Cy7-A :: NKp46
Comp-FITC-A :: NK11
Comp-Qdot 605-A :: CD19
Comp-Pacific Blue-A :: CD3
Comp-PerCP-Cy5-5-A :: CD4
Comp-Alexa Fluor 700-A :: CD8
Comp-PE-A :: FOXP3
Comp-APC-A :: CD25

b.

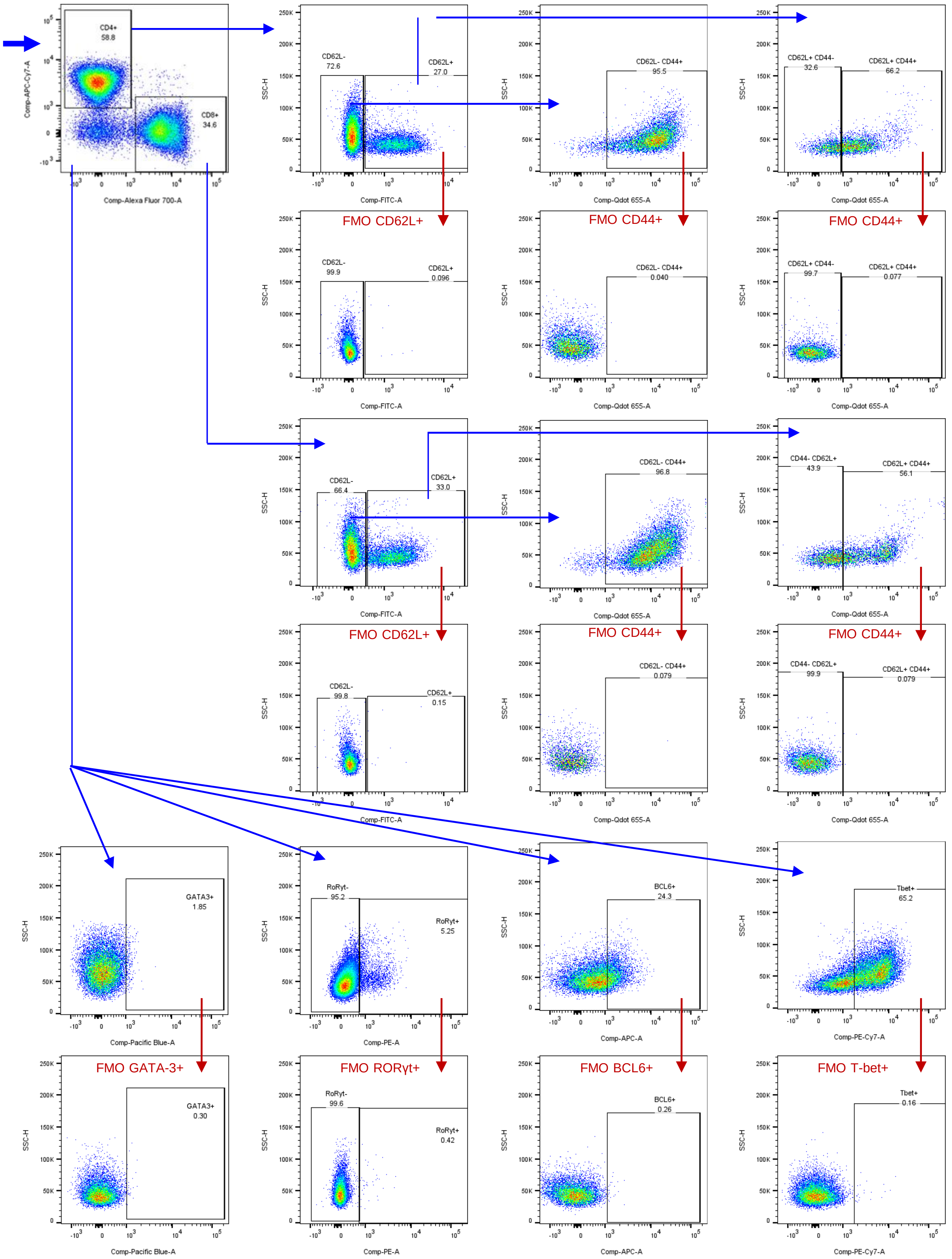

CD4+
58.8
CD8+
34.6
Comp-APC-Cy7-A
Comp-Alexa Fluor 700-A
CD62L-
72.6
CD62L+
27.0
CD62L- CD44+
95.5
CD62L+ CD44-
32.6
CD62L+ CD44+
66.2
FMO CD62L+
CD62L-
99.9
CD62L+
0.096
FMO CD44+
CD62L- CD44+
0.040
FMO CD44+
CD62L+ CD44-
99.7
CD62L+ CD44+
0.077
CD62L-
66.4
CD62L+
33.0
CD62L- CD44+
96.8
CD44- CD62L+
43.9
CD62L+ CD44+
56.1
FMO CD62L+
CD62L-
99.8
CD62L+
0.15
FMO CD44+
CD62L- CD44+
0.079
FMO CD44+
CD44- CD62L+
99.9
CD62L+ CD44+
0.079
GATA3+
1.85
RoRyt-
95.2
RoRyt+
5.25
BCL6+
24.3
Tbet+
65.2
FMO GATA-3+
GATA3+
0.30
FMO RORγt+
RoRyt-
99.6
RoRyt+
0.42
FMO BCL6+
BCL6+
0.26
FMO T-bet+
Tbet+
0.16
SSC-H
Comp-FITC-A
Comp-Qdot 655-A
Comp-Pacific Blue-A
Comp-PE-A
Comp-APC-A
Comp-PE-Cy7-A

c.

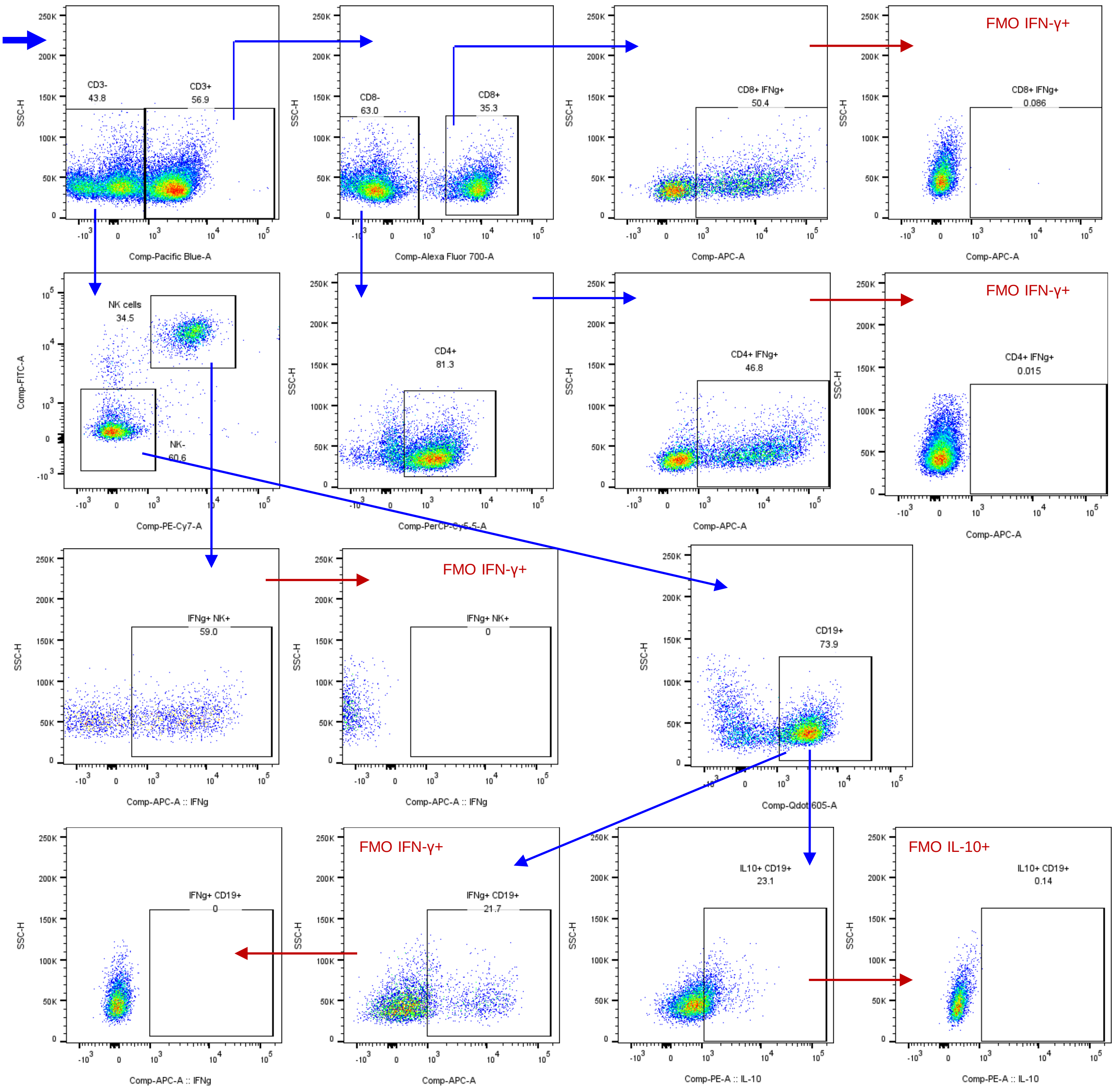

CD3-
43.8
CD3+
56.9
Comp-Pacific Blue-A
CD8-
63.0
CD8+
35.3
Comp-Alexa Fluor 700-A
CD8+ IFNg+
50.4
Comp-APC-A
FMO IFN-γ+
CD8+ IFNg+
0.086
Comp-APC-A
NK cells
34.5
NK-
60.6
Comp-FITC-A
Comp-PE-Cy7-A
CD4+
81.3
Comp-PerCP-Cy5-5-A
CD4+ IFNg+
46.8
Comp-APC-A
FMO IFN-γ+
CD4+ IFNg+
0.015
Comp-APC-A
IFNg+ NK+
59.0
Comp-APC-A :: IFNg
FMO IFN-γ+
IFNg+ NK+
0
Comp-APC-A :: IFNg
CD19+
73.9
Comp-Qdot 605-A
IFNg+ CD19+
0
Comp-APC-A :: IFNg
FMO IFN-γ+
IFNg+ CD19+
21.7
Comp-APC-A
IL10+ CD19+
23.1
Comp-PE-A :: IL-10
FMO IL-10+
IL10+ CD19+
0.14
Comp-PE-A :: IL-10
SSC-H

**Figure S5. Two-dimensional gating strategies for identification of lymphoid cell subsets by flow cytometry.** As described in the Methods and shown in Fig. 4b–t, FCG mice were euthanized at 4 and 12 wpi. Single-cell suspensions from lungs were prepared, stained with the indicated antibodies, and analyzed by multicolor flow cytometry. **(a)**. Gating strategy for identification of total $CD4^+$ and $CD8^+$ T cells, NK cells, B cells, and regulatory T cells (Tregs). **(b)**. Gating strategy for identification of total central memory (TCM), effector memory (TEM), and naïve subsets within $CD4^+$ and $CD8^+$ T cell populations. Also shown are gating strategies for transcription factor–defined $CD4^+$ T cell subsets, including $GATA3^+$ (Th2), $ROR\gamma t^+$ (Th17), Bcl-$6^+$ (Tfh), and T-$bet^+$ (Th1) cells. At 4 and 12 wpi, lung single-cell suspensions were stimulated for 4 h (as detailed in the Methods), and IFN-γ and IL-10 expression across lymphoid subsets was assessed by multicolor flow cytometry. **(c).** Gating strategy for identification of IFN-$\gamma^+$ $CD4^+$ T cells, IFN-$\gamma^+$ $CD8^+$ T cells, IFN-$\gamma^+$ NK cells, IFN-$\gamma^+$ B effector 1 cells, and IL-$10^+$ B cells. All antibodies were titrated to determine optimal concentrations that maximized specificity while minimizing spectral spillover. Gating strategies were established using single-stained controls and fluorescence-minus-one (FMO) controls. Gates are shown in blue/black, and corresponding FMO controls are indicated in red.

FigS6.

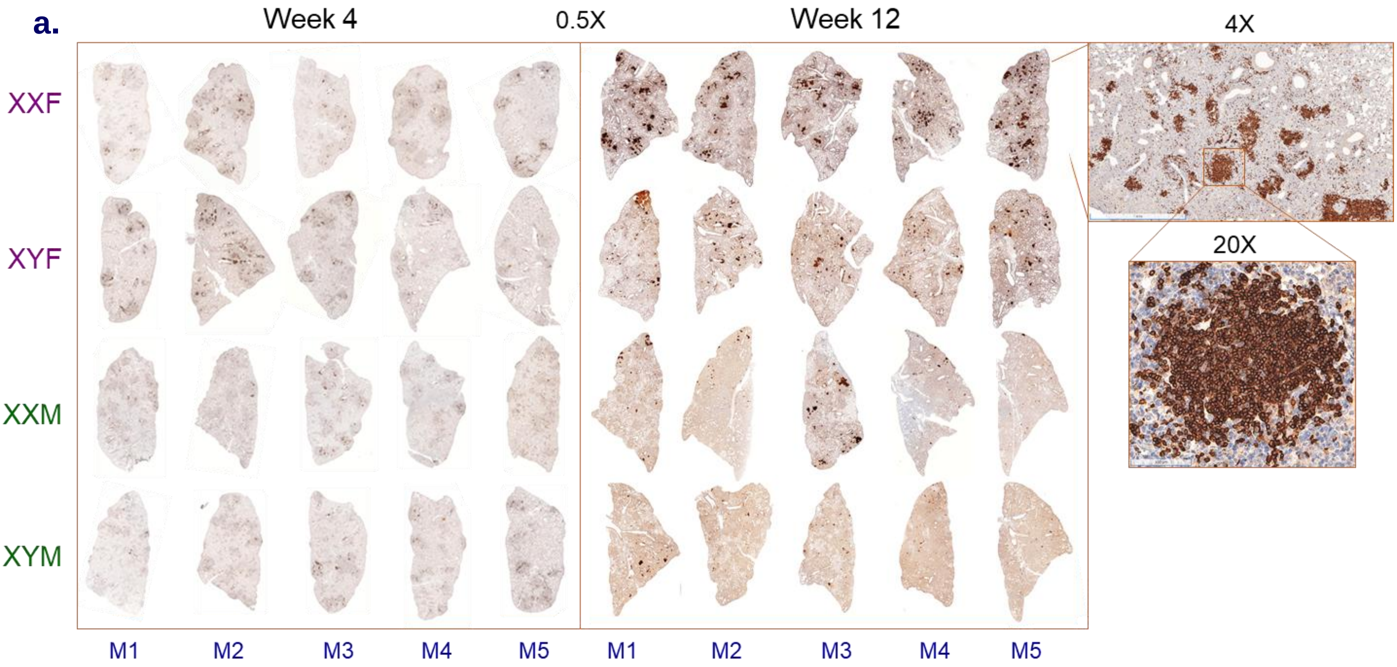


**Figure S6. Enhanced B cell follicle formation and activation in gonadal females. (a).** As shown in Fig. 5a, formalin-fixed lung tissues collected at 4 and 12 weeks post-infection were sectioned (same samples as in Fig. S2a) and stained with anti-B220/CD45R antibodies. Representative B220/CD45R-stained lung sections are shown at 0.5× magnification. Insets highlight B220/CD45R$^{+}$ areas within localized granulomas at 4× and 20× magnification. **(b).** Gating strategy for identification of total B cells (BC), B-1a BC, B-1b BC, B2 BC, follicular BC, germinal center (GC) BC, memory BC, and plasmablasts by flow cytometry. All antibodies were titrated to determine optimal concentrations that maximized specificity while minimizing spectral spillover. Gating strategies were established using single-stained controls and fluorescence-minus-one (FMO) controls. Gates are shown in blue/black, and corresponding FMO controls are indicated in red.

**b.**

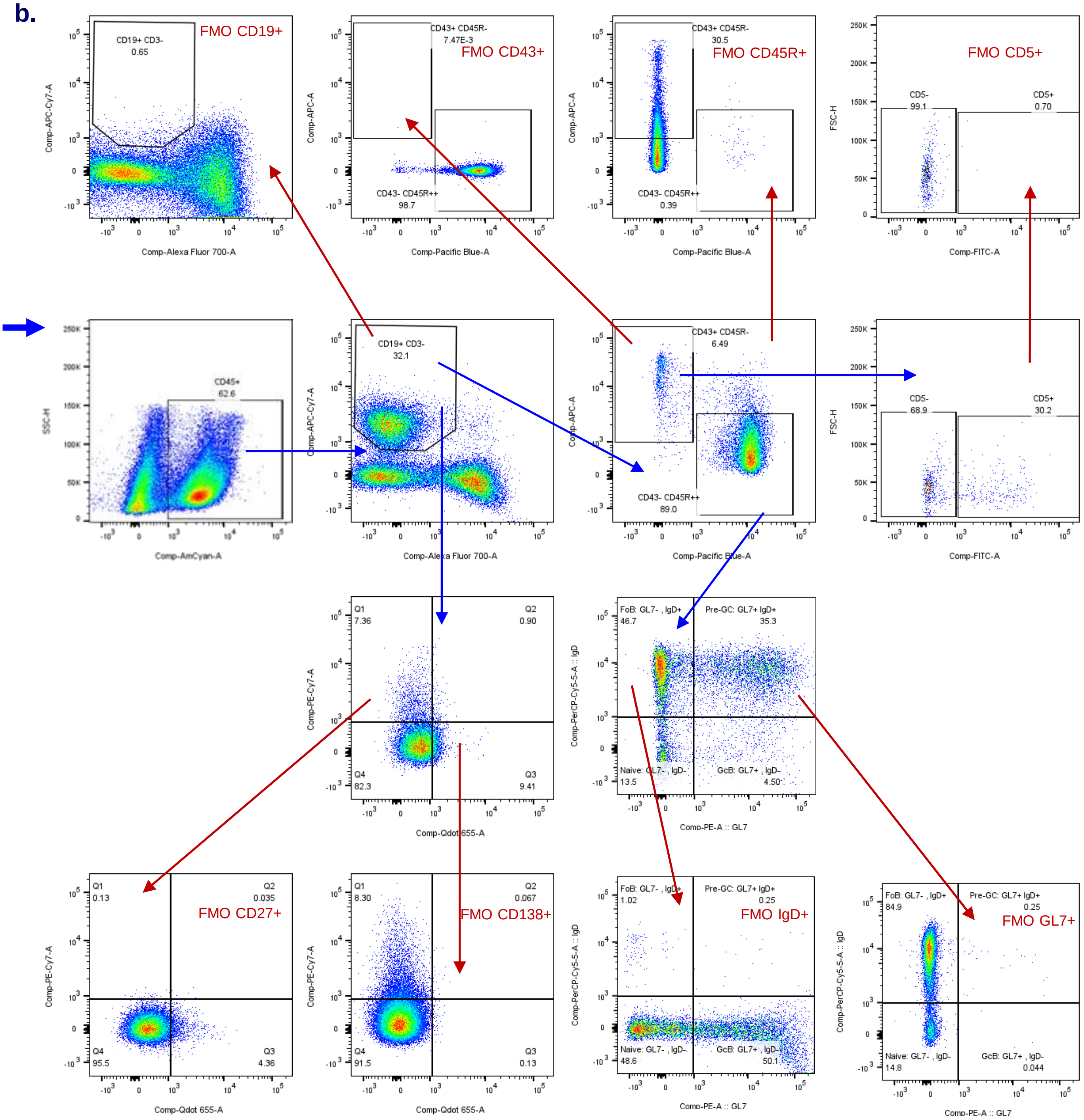
FMO CD19+
CD19+ CD3-
0.65
Comp-APC-Cy7-A
Comp-Alexa Fluor 700-A
FMO CD43+
CD43+ CD45R-
7.47E-3
CD43- CD45R++
98.7
Comp-APC-A
Comp-Pacific Blue-A
FMO CD45R+
CD43+ CD45R-
30.5
CD43- CD45R++
0.39
FMO CD5+
CD5-
99.1
CD5+
0.70
FSC-H
Comp-FITC-A
CD45+
62.6
SSC-H
Comp-AmCyan-A
CD19+ CD3-
32.1
CD43+ CD45R-
6.49
CD43- CD45R++
89.0
CD5-
68.9
CD5+
30.2
Q1
7.36
Q2
0.90
Q4
82.3
Q3
9.41
Comp-PE-Cy7-A
Comp-Qdot 655-A
FoB: GL7- , IgD+
46.7
Pre-GC: GL7+ IgD+
35.3
Naive: GL7- , IgD-
13.5
GcB: GL7+ , IgD-
4.50
Comp-PerCP-Cy5-5-A :: IgD
Comp-PE-A :: GL7
FMO CD27+
Q1
0.13
Q2
0.035
Q4
95.5
Q3
4.36
FMO CD138+
Q1
8.30
Q2
0.067
Q4
91.5
Q3
0.13
FMO IgD+
FoB: GL7- , IgD+
1.02
Pre-GC: GL7+ IgD+
0.25
Naive: GL7- , IgD-
48.6
GcB: GL7+ , IgD-
50.1
FMO GL7+
FoB: GL7- , IgD+
84.9
Pre-GC: GL7+ IgD+
0.25
Naive: GL7- , IgD-
14.8
GcB: GL7+ , IgD-
0.044

### FigS7.

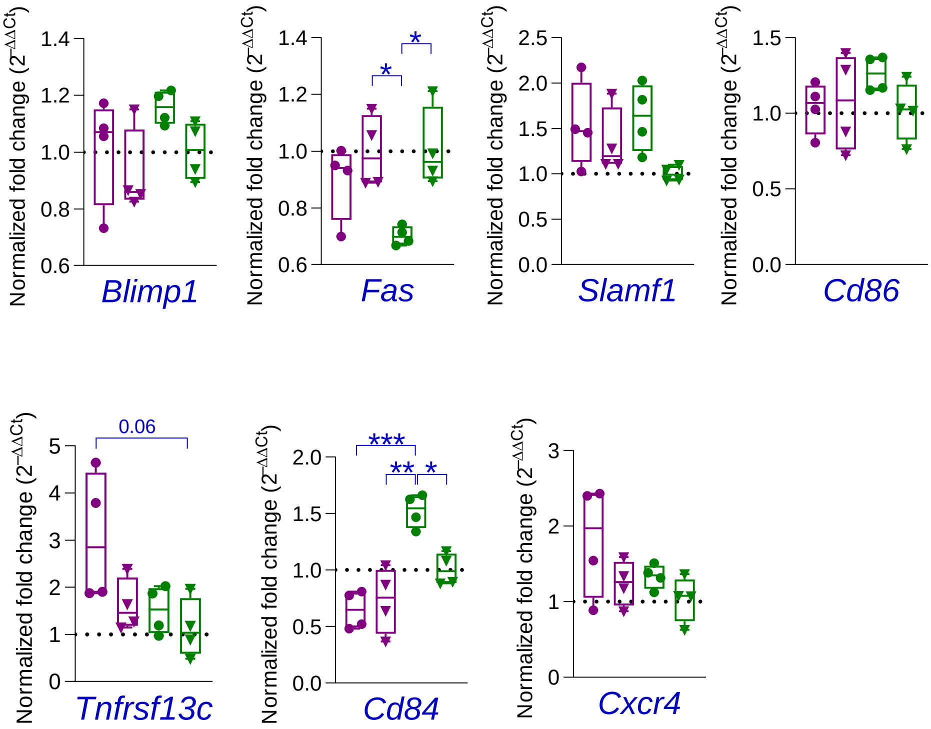


**Figure S7. Genes involved in Tfh-mediated B cell help did not exhibit sex-biased expression.** qPCR analysis of various genes involved in BCF formation, regulation, and maintenance. 18S rRNA normalized gene expression of various genes measured by qPCR at 12 wpi (n=4). Data shown as fold change relative to XYM (dotted line). Fold change calculated by $2^{-\Delta\Delta Ct}$ method. Each dot represents an individual mouse and is the average of duplicates. Data are presented as mean ± SEM values. Statistical significance was calculated using two-way ANOVA. *P < 0.05, ** P < 0.01, *** P < 0.001 and ns for no significance.
